# Cryo-EM structures of human zinc transporter ZnT7 reveal the mechanism of Zn^2+^ uptake into the Golgi apparatus

**DOI:** 10.1101/2022.03.23.485435

**Authors:** Bui Ba Han, Satoshi Watanabe, Norimichi Nomura, Kehong Liu, Tomoko Uemura, Michio Inoue, Akihisa Tsutsumi, Hiroyuki Fujita, Kengo Kinoshita, So Iwata, Masahide Kikkawa, Kenji Inaba

## Abstract

Zinc ions (Zn^2+^) are vital to most cells, with the intracellular concentrations of Zn^2+^ being tightly regulated by multiple zinc transporters located at the plasma and organelle membranes. We herein present the 2.8-2.9 Å-resolution cryo-EM structures of a Golgi-localized human Zn^2+^/H^+^ antiporter ZnT7 (hZnT7) in its outward- and inward-facing forms. Cryo-EM analyses showed that hZnT7 exists as a homodimer via tight interactions in both the cytosolic and transmembrane (TM) regions of two protomers, each of which contains a single Zn^2+^-binding site in its TM domain. hZnT7 undergoes a TM-helix rearrangement to create a negatively charged cytosolic cavity for Zn^2+^ entry in the inward-facing form and a widened luminal cavity for Zn^2+^ release in the outward-facing form. An exceptionally long cytosolic histidine-rich loop characteristic of hZnT7 can bind at least two Zn^2+^ ions, likely facilitating Zn^2+^ recruitment from the cytosol. Unique mechanisms of hZnT7-mediated Zn^2+^ uptake into the Golgi are proposed.

## Introduction

Zinc ions (Zn^2+^) are an essential trace element that plays an important role in the structure and function of a variety of proteins, acting as either a cofactor essential for enzymatic reactions or as a structural element stabilizing protein folding. Nearly 10% of the mammalian proteome is known to bind Zn^2+1^. Cellular zinc homeostasis involves the opposing actions of two families of zinc transporters, SLC30 (ZnTs) and SLC39 (ZIPs)^2^. The SLC30 family proteins transport Zn^2+^ from the cytosol to the extracellular space or intracellular compartments, thereby reducing cytosolic Zn^2+^ concentrations. In human cells, the SLC30 (ZnTs) family consists of 10 homologues, named ZnT1 to ZnT10, which are largely responsible for the dynamics of intracellular and extracellular Zn^2+3, 4^. One of these proteins, human ZnT7 (hZnT7), has been reported to localize in the Golgi membrane^5–8^ and transport Zn^2+^ from the cytosol into the Golgi lumen for incorporation into newly synthesized zinc-enzymes^3, 4^. Due to the presence of ZnT7 and other Golgi-resident ZnTs, the labile Zn^2+^ concentration in the Golgi is maintained at around 25 nM or higher^9^. ZnT7 functions as a Zn^2+^/H^+^ antiporter and uses proton motive force to transport Zn^2+^ from the cytosol to the Golgi lumen. Our latest study showed that ZnT7 localizes on the proximal side of the Golgi and contributes to the regulation of the intracellular localization and traffic of ERp44, an ER-Golgi cycling chaperone^10^.

Physiologically, ZnT7 plays essential roles in dietary zinc absorption and regulation of body adiposity^11^. Decrease of cellular Zn^2+^ in the epithelium of the prostate was shown to be involved in the development of prostate cancer in mice, with apoptosis being prevented in TRAMP/ZnT7(-/-) mice^12^. Therefore, a null mutation in the Znt7 gene accelerated prostate tumor formation in mice^12^. In addition, a deficiency in ZnT7 reduced lipid synthesis in adipocytes by inhibiting insulin-dependent Akt activation and glucose uptake^13^.

Despite studies of the physiology of ZnT7, its structure and Zn^2+^ transport mechanism remain to be elucidated. Structures have been determined for several zinc transporters, including *Escherichia coli* YiiP (EcYiiP)^14, 15^, *Shewanella oneidensis* YiiP (soYiiP)^16–, 18^, and human ZnT8 (hZnT8)^19^, all of which belong to the SLC30 family. Despite being at non-atomic resolution, cryo-EM analysis of hZnT8, a Zn^2+^/H^+^ antiporter localized to the insulin secretory granules of pancreatic β cells, revealed its overall architecture and Zn^2+^-binding sites^19^. Amino acid sequence alignment (Fig. 1) suggests significant differences between hZnT7 and hZnT8, especially in their overall architecture, length of their cytosolic histidine-rich loops (His-loop), and numbers of Zn^2+^-binding sites. Because of its physiological importance, hZnT7 is a particular target of structural and mechanistic studies.

**Fig. 1.**
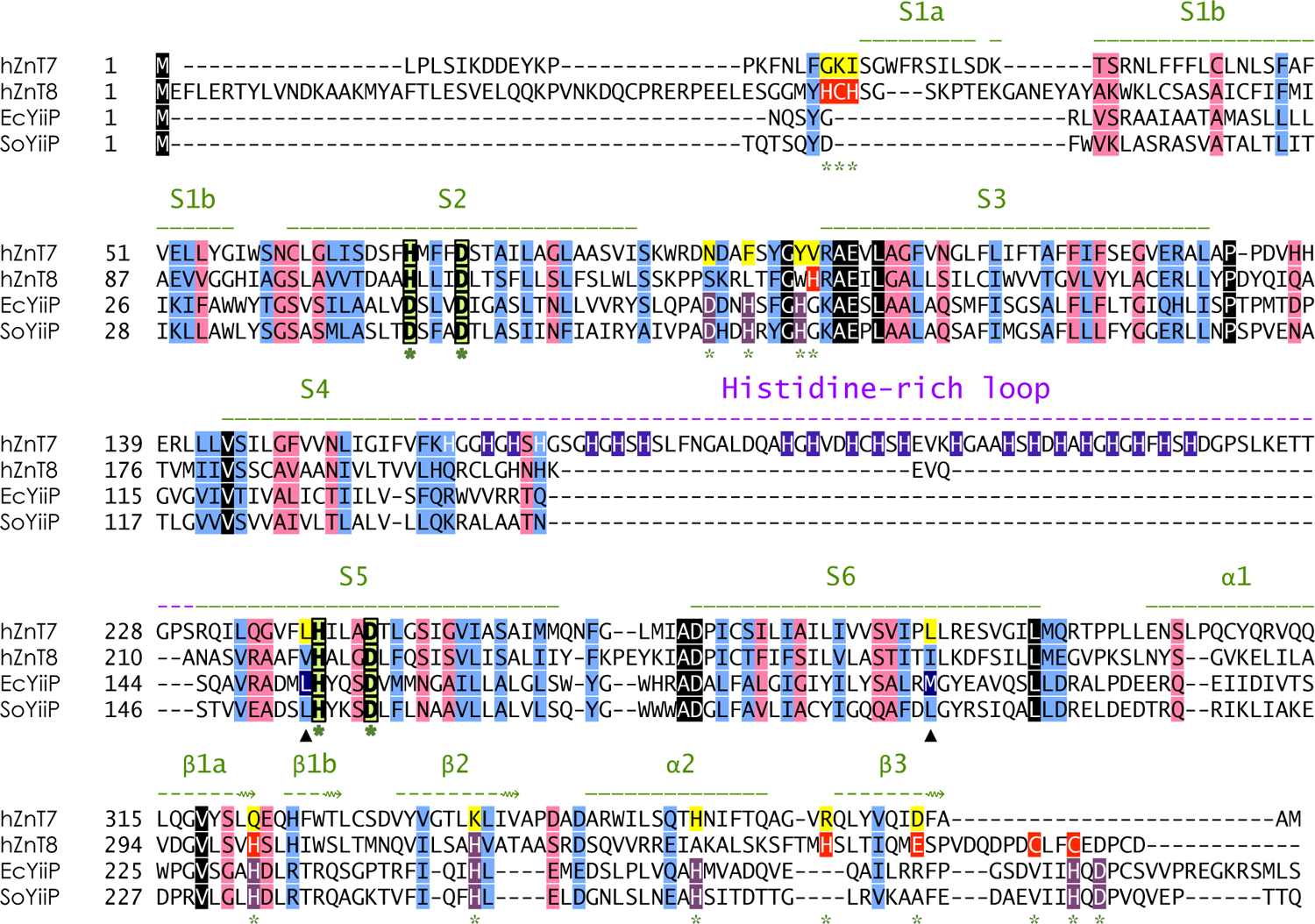
Sequence alignment of representative ZnT-family members of known structure. Amino acid sequences of human ZnT7 (hZnT7) (UniProtKB - Q8NEW0) and ZnT8 (hZnT8) (UniProtKB - Q8IWU4), *Escherichia coli* YiiP (EcYiiP) (UniProtKB - P69380), and *Shewanella oneidensis* YiiP (SoYiiP) (UniProtKB - Q8E919) were aligned. Highly conserved, similar, and weakly conserved residues are colored black, light-blue, and rose, respectively. The α-helix and β-strand elements of hZnT7 are indicated by solid and broken green lines above the amino acid sequence, respectively. The histidine-rich loop is indicated by a broken pink line above the sequence. HDHD-motif residues present in the transmembrane domain are indicated by light green boxes; Zn^2+^-binding residues present in the cytosolic domain of hZnT8 and Ec/SoYiiP are indicated by red and violet boxes, respectively; and histidine residues in the His-loop of ZnT7 are indicated by blue boxes. Green stars indicate residues involved in Zn^2+^ binding. S, transmembrane helix; α and β, alpha-helix and beta-strand in the cytosolic domain, respectively.

The present study describes the assessment of high-resolution cryogenic electron microscopy (cryo-EM) structures of hZnT7 in its Zn^2+^-bound and -unbound states. A monoclonal antibody Fab fragment that specifically and tightly binds to native-state hZnT7 was prepared, and the structures of the hZnT7-Fab complex were determined by single-particle cryo-EM analysis. Consequently, the structures of the Zn^2+^-unbound outward-facing (OF) and inward-facing (IF) forms, and Zn^2+^-bound OF form of hZnT7 were determined at 2.8, 3.4, and 2.9 Å resolutions, respectively. Determinations of these structures provided essential insight into mechanisms of hZnT7-mediated Zn^2+^ transport into the Golgi. Moreover, the roles of the His-loop were characterized by preparing a His-loop deletion mutant of hZnT7 and comparing its structure and Zn^2+^-binding property with those of wild-type hZnT7. These findings enable comparisons of the structural and mechanistic features of hZnT7 with those of other Zn^2+^ transporters of known structure, and have physiological implications for Zn^2+^ homeostasis in the Golgi apparatus.

## Results and Discussion

### Overall structure of hZnT7 in the Zn^2+^-unbound state

We employed single-particle cryo-electron microscopy (cryo-EM) analysis to determine the structure of the 74-kDa hZnT7 homodimer. After data processing of the repeated 2D- and 3D-classifications, a density map of the hZnT7 homodimer was obtained at ∼12 Å resolution (data not shown), which did not allow *de novo* model building. This problem was overcome by preparing monoclonal antibody Fab fragments against recombinant hZnT7 to increase the molecular size of the latter and thereby generate higher resolution cryo-EM maps (see “<u>Methods</u>” section). After thorough screening by ELISA and size-exclusion chromatography (SEC), four candidate monoclonal antibodies were selected, their Fab fragments (Fab#1, #3, #4, and #5) were generated, and four types of hZnT7-Fab complexes were prepared (**Supplementary Fig. 1**). Two of these complexes, hZnT7-Fab#1 and hZnT7-Fab#3, yielded high quality two-dimensional class averages by negative-stain EM measurements, showing that a Fab fragment symmetrically bound to each subunit of the hZnT7 homodimer (**Supplementary Fig. 2a**). The hZnT7 proteins in complex with Fab#1 and Fab#3 were analyzed by Talos Arctica cryo-transmission electron microscopy (TEM), yielding 5.0 Å- and 7.7 Å-resolution density maps, respectively (**Supplementary Fig. 2b**). These results indicated that Fab#1 was the optimal binder of hZnT7 for cryo-EM analysis. The structure of the hZnT7-Fab#1 complex was therefore assessed by high-end Titan Krios cryo-TEM with a Gatan K3 BioQuantum direct electron detector.

A cryo-EM structure of the hZnT7-Fab#1 complex at a nominal resolution of 2.8 Å was determined (**Fig. 2a, Supplementary Figs. 3-4**, and **Table 1**). Four rounds of 3D classification generated a major class of density map (class 2), which revealed a “*mushroom*”-shaped dimeric architecture of hZnT7 (**Fig. 2b** and **2c**) and the mode of interaction between hZnT7 and the Fab fragment (**Supplementary Fig. 6**). The Fab fragment specifically bound to the α1-β1a and β2-α2 loops, which are both located at the end of the hZnT7 cytosolic domain (CTD). No direct interactions were made between the Fab fragment and the transmembrane domain (TMD) or the intermediate regions of the TMD and CTD of hZnT7. It seems unlikely that Fab binding considerably affects the overall fold and transmembrane (TM) helix arrangement of hZnT7. (**Fig. 2b**). The TMD of the hZnT7 protomer was found to contain six transmembrane helices (S1 to S6), with both the amino and carboxyl termini exposed to the cytosol. This protomer could be divided into two subdomains, a “*scaffold*” domain composed of S1 to S3 and S6, and a “*helix-pair*” domain composed of S4 and S5. These two subdomains formed a Zn^2+^ transport cavity (**Fig. 2b**), in which S2 and S5 formed a Zn^2+^-binding HDHD motif composed of His^70^ (S2), Asp^74^ (S2), His^240^ (S5), and Asp^244^ (S5) at their midway points. Careful examination of the density map, however, indicated that no Zn^2+^ was bound to hZnT7 (**Fig. 2d**), suggesting that Zn^2+^ had been released from hZnT7 during purification procedures using a Zn^2+^-free buffer.

**Fig. 2.**
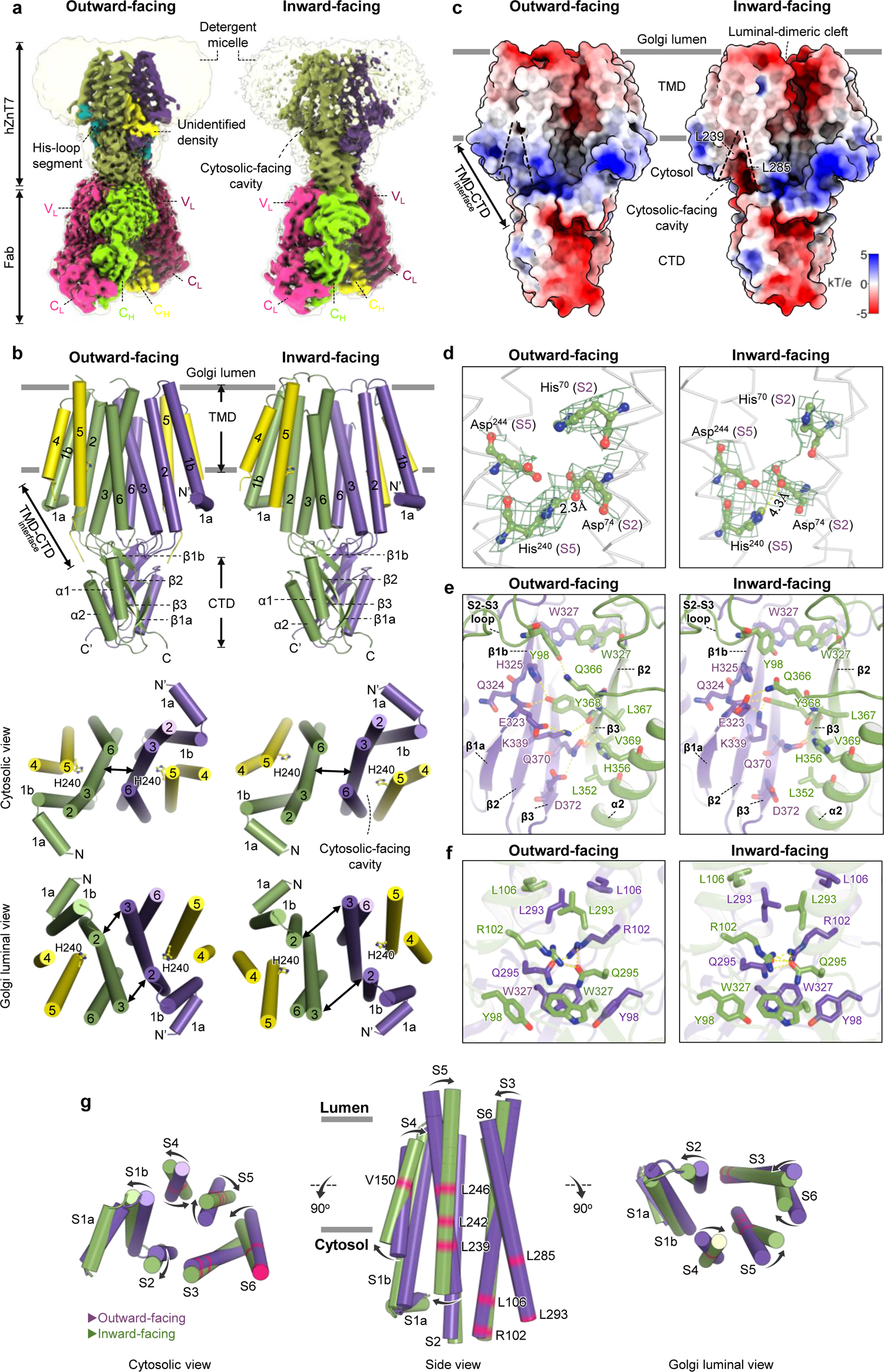
Cryo-EM structures of Zn^2+^-unbound outward-facing (OF) and inward-facing (IF) forms of hZnT7. (**a**) Side views of outward-facing (left) and inward-facing (right) forms of hZnT7 in complex with -Fab#1, with density maps shown in surface representations at a contour level of 0.5σ. The density of detergent micelles is shown in the surface representation as a transparent envelope at a contour level of 0.25σ. (**b**) Cylinder representation of the hZnT7 dimer in the same orientation as in (a) (top), and cytosolic (middle) and luminal (bottom) views of the transmembrane domain of hZnT7, with the cytosolic domain and TM loops removed for clarity. The membrane boundaries were determined using the OPM server (http://opm.phar.umich.edu/server.php) and are shown with gray lines (top). S1 to S3 and S6 (shown in green or violet) form the scaffold domain, while S4 and S5 (shown in yellow) form the helix-pair domain. TMD, transmembrane domain; CTD, cytosolic domain. (**c**) Electrostatic potential surface map of outward-facing and inward-facing forms of hZnT7 at pH 7.5. Black dashed lines indicate the cytosolic-facing cavity present at the TMD-CTD interface. Gray lines indicate boundaries of the membrane. Surface colors represent Coulombic potential (red, negative; white, neutral; blue, positive). T, temperature; k, Boltzmann constant; e, charge of an electron. (**d**) Zn^2+^-binding sites in the transmembrane domain of outward-facing (left) and inward-facing (right) forms of hZnT7. Residues forming the HDHD motif are represented by sticks, while the density of these residues is shown by green mesh at contour levels of 0.4σ for outward-facing and 7σ for inward-facing forms. (**e**) CTD interactions at the dimer interface in outward-facing (left) and inward-facing (right) forms of hZnT7. Yellow dotted lines indicate contacts between polar residues. (**f**) The TMD-CTD interface in outward-facing (left) and inward-facing (right) forms of hZnT7. (**g**) Superposition of the TMDs of outward-facing (purple) and inward-facing (green) forms of hZnT7. Black arrows indicate the movement of TM helices during the conversion from the outward-facing to the inward-facing form.

**Fig. 3.**
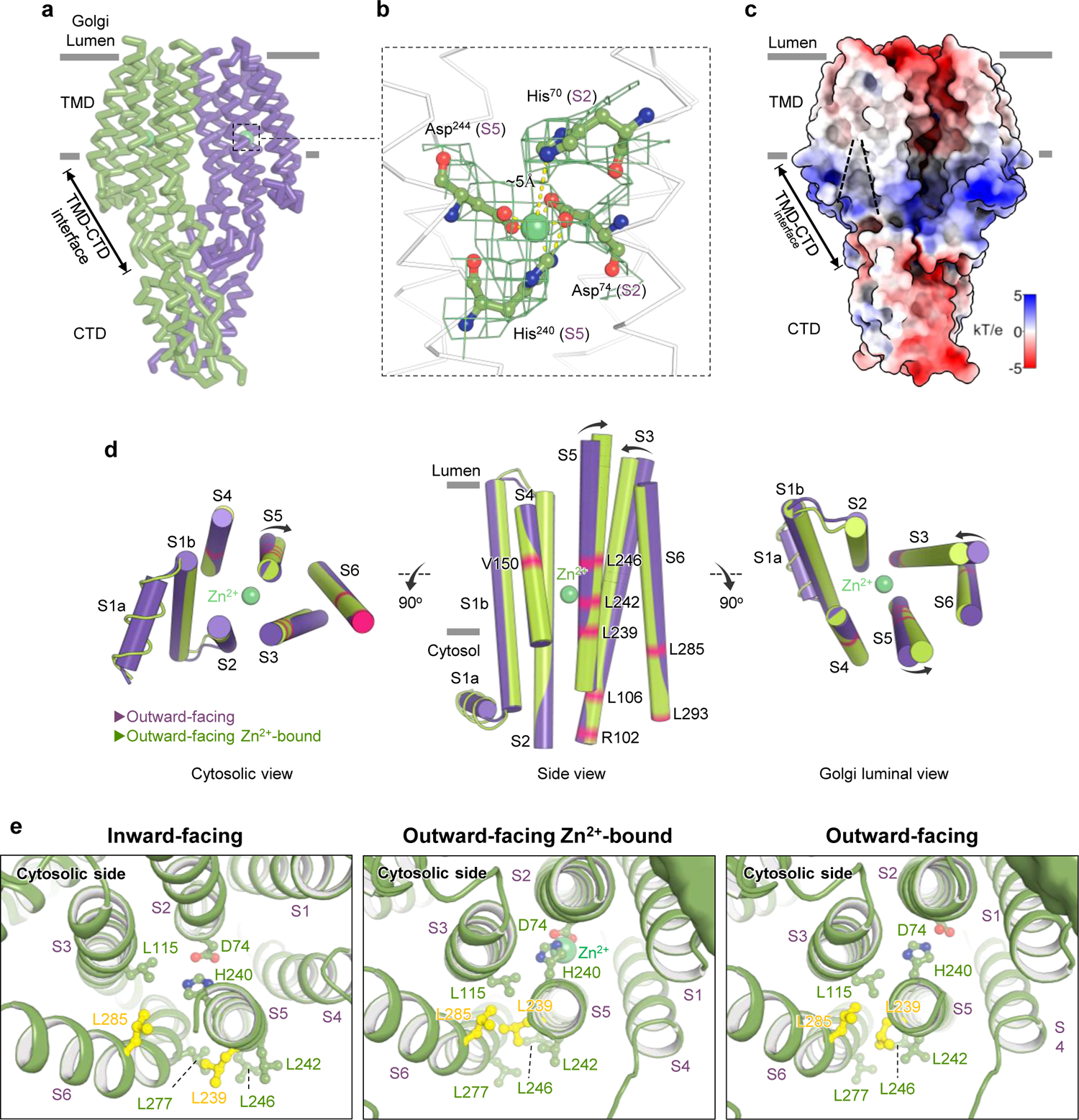
Cryo-EM structures of the Zn^2+^-bound outward-facing form of hZnT7. (**a**) Ribbon diagram showing the overall structure of the outward-facing form of Zn^2+^-bound hZnT7. Bound Zn^2+^ ions are represented by green spheres. (**b**) The Zn^2+^-binding site viewed from the cytosolic side. Residues forming the HDHD motif are represented by ball-and-sticks, while the density of these residues is shown by green mesh at a contour level of 0.4σ. (**c**) Electrostatic potential surface map of the outward-facing form of Zn^2+^-bound hZnT7 at pH 7.5. The orientation is the same as in (a). Black dashed lines indicate the cytosolic-facing cavity present at the TMD-CTD interface. Gray lines indicate boundaries of the membrane. Surface colors represent Coulombic potential (red, negative; white, neutral; blue, positive). T, temperature; k, Boltzmann constant; e, charge of an electron. (**d**) Superposition of the TM helices of the outward-facing forms of Zn^2+^-bound (light green) and -unbound (violet) hZnT7. Bound Zn^2+^ is represented by a green sphere. Black arrows indicate the movement of TM helices induced by Zn^2+^ binding. (**e**) The Leu^239^ (S5)-Leu^285^ (S6) gate that likely controls the opening/closure of the cytosolic-facing cavity viewed from the cytosolic side. Leu^293^ (S5) and Leu^285^ (S6) are represented by yellow sticks. The cytosolic domain is omitted for clarity.

**Fig. 4.**
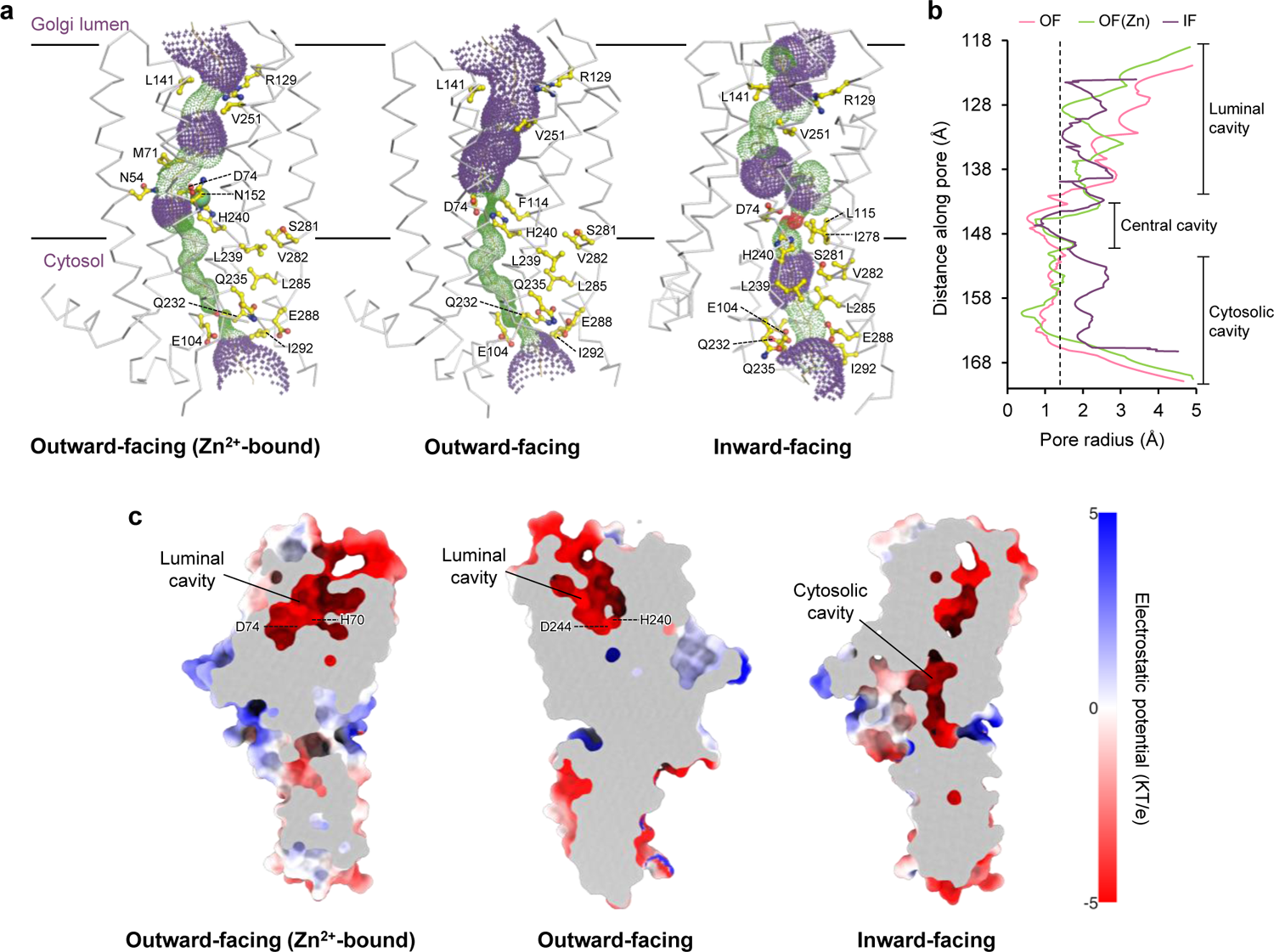
Zinc ion permeation pore of hZnT7. (**a**) Pore profile of a hZnT7 protomer in outward-facing forms of the Zn^2+^-bound (left) and -unbound (middle) states, and in the inward-facing form of the Zn^2+^-unbound state (right). For clarity, only the TMD is highlighted. Red, green, and purple mesh colors correspond to pore radii of <1.0 Å, 1.0-2.3 Å, and >2.3Å, respectively. The protein backbones are represented by ribbons, and the zinc ion is represented by a green sphere. Black lines indicate boundaries of the membrane. (**b**) Pore radius of the hZnT7 protomer calculated using a HOLE program. Colors correspond to the outward-facing forms of Zn^2+^-bound (OFZ, green) and -unbound (OF, red) states, and the inward-facing form of the Zn^2+^-unbound state (IF, violet). The vertical broken line indicates the radius of a water molecule, 1.4 Å. (**c**) Coronal section of the electrostatic potential surface map of the hZnT7 protomer. Surface colors indicate Coulombic potentials (red, negative; white, neutral; blue, positive). T, temperature; k, Boltzmann constant; e, charge of an electron.

**Table 1.**
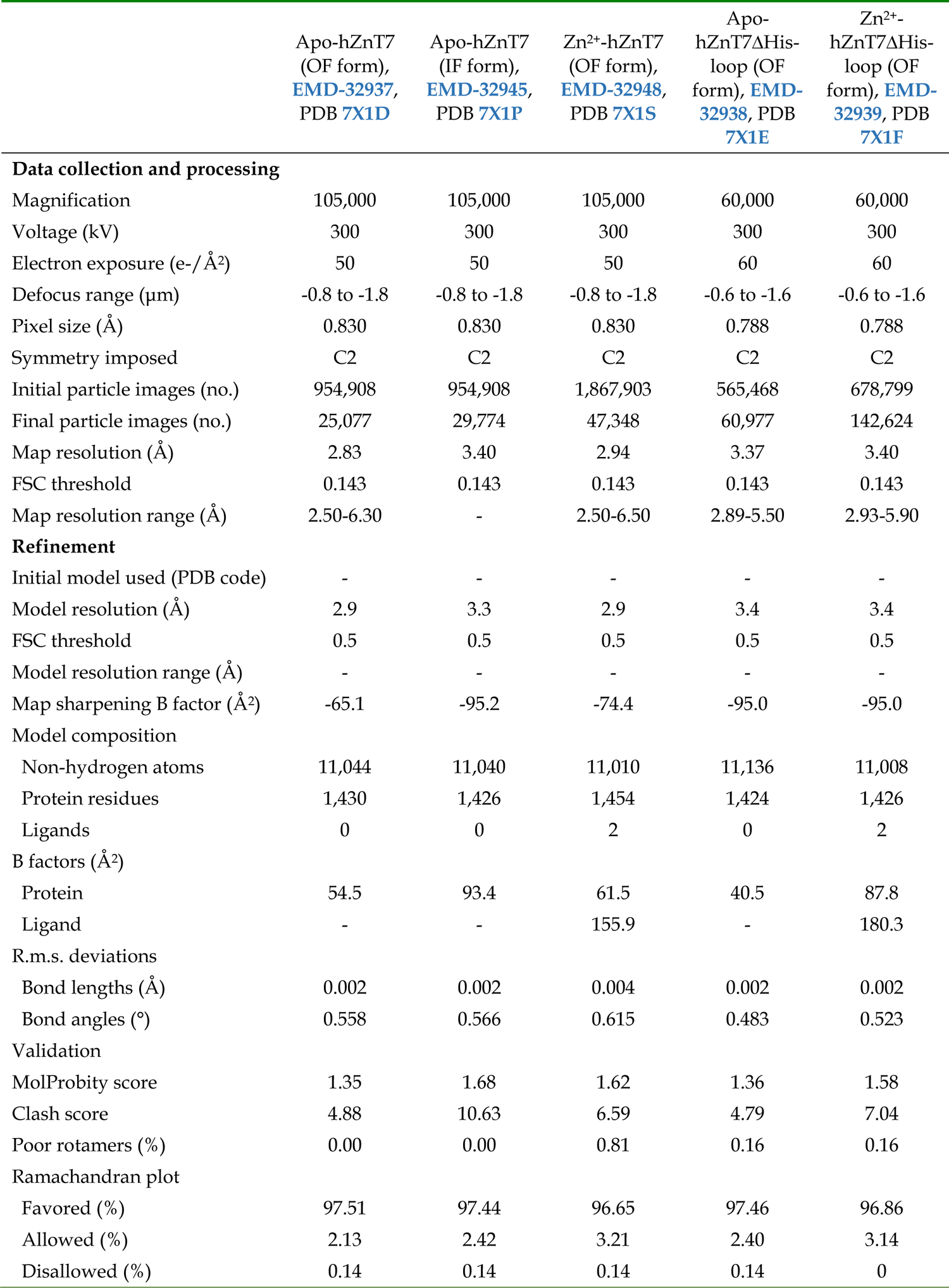
Cryo-EM data collection, refinement, and validation statistics.

|  | Apo-hZnT7<br>(OF form),<br>EMD-32937,<br>PDB 7X1D | Apo-hZnT7<br>(IF form),<br>EMD-32945,<br>PDB 7X1P | Zn<sup>2+</sup>-hZnT7<br>(OF form),<br>EMD-32948,<br>PDB 7X1S | Apo-<br>hZnT7ΔHis-<br>loop (OF<br>form), EMD-<br>32938, PDB<br>7X1E | Zn<sup>2+</sup>-<br>hZnT7ΔHis-<br>loop (OF<br>form), EMD-<br>32939, PDB<br>7X1F |
| --- | --- | --- | --- | --- | --- |
| **Data collection and processing** |  |  |  |  |  |
| Magnification | 105,000 | 105,000 | 105,000 | 60,000 | 60,000 |
| Voltage (kV) | 300 | 300 | 300 | 300 | 300 |
| Electron exposure (e-/Å<sup>2</sup>) | 50 | 50 | 50 | 60 | 60 |
| Defocus range (μm) | -0.8 to -1.8 | -0.8 to -1.8 | -0.8 to -1.8 | -0.6 to -1.6 | -0.6 to -1.6 |
| Pixel size (Å) | 0.830 | 0.830 | 0.830 | 0.788 | 0.788 |
| Symmetry imposed | C2 | C2 | C2 | C2 | C2 |
| Initial particle images (no.) | 954,908 | 954,908 | 1,867,903 | 565,468 | 678,799 |
| Final particle images (no.) | 25,077 | 29,774 | 47,348 | 60,977 | 142,624 |
| Map resolution (Å) | 2.83 | 3.40 | 2.94 | 3.37 | 3.40 |
| FSC threshold | 0.143 | 0.143 | 0.143 | 0.143 | 0.143 |
| Map resolution range (Å) | 2.50-6.30 | - | 2.50-6.50 | 2.89-5.50 | 2.93-5.90 |
| **Refinement** |  |  |  |  |  |
| Initial model used (PDB code) | - | - | - | - | - |
| Model resolution (Å) | 2.9 | 3.3 | 2.9 | 3.4 | 3.4 |
| FSC threshold | 0.5 | 0.5 | 0.5 | 0.5 | 0.5 |
| Model resolution range (Å) | - | - | - | - | - |
| Map sharpening B factor (Å<sup>2</sup>) | -65.1 | -95.2 | -74.4 | -95.0 | -95.0 |
| **Model composition** |  |  |  |  |  |
| Non-hydrogen atoms | 11,044 | 11,040 | 11,010 | 11,136 | 11,008 |
| Protein residues | 1,430 | 1,426 | 1,454 | 1,424 | 1,426 |
| Ligands | 0 | 0 | 2 | 0 | 2 |
| **B factors (Å<sup>2</sup>)** |  |  |  |  |  |
| Protein | 54.5 | 93.4 | 61.5 | 40.5 | 87.8 |
| Ligand | - | - | 155.9 | - | 180.3 |
| **R.m.s. deviations** |  |  |  |  |  |
| Bond lengths (Å) | 0.002 | 0.002 | 0.004 | 0.002 | 0.002 |
| Bond angles (°) | 0.558 | 0.566 | 0.615 | 0.483 | 0.523 |
| **Validation** |  |  |  |  |  |
| MolProbity score | 1.35 | 1.68 | 1.62 | 1.36 | 1.58 |
| Clash score | 4.88 | 10.63 | 6.59 | 4.79 | 7.04 |
| Poor rotamers (%) | 0.00 | 0.00 | 0.81 | 0.16 | 0.16 |
| **Ramachandran plot** |  |  |  |  |  |
| Favored (%) | 97.51 | 97.44 | 96.65 | 97.46 | 96.86 |
| Allowed (%) | 2.13 | 2.42 | 3.21 | 2.40 | 3.14 |
| Disallowed (%) | 0.14 | 0.14 | 0.14 | 0.14 | 0 |

The cavity formed by S2, S3, and S5 was significantly wider at the Golgi luminal side than at the cytosolic side (**Fig. 2b** and **2g**), suggesting that this major-class structure represents the OF form of hZnT7. In addition to binding Zn^2+^, the TMD plays an essential role for dimerization, in that S2 from one protomer makes hydrophobic contact with S3 from another protomer, forming a tight dimer interface (**Fig. 2b, Supplementary Fig. 7a**, and **Supplementary Fig. 8**). Additionally, S3 from one protomer interacts with S3 from another promoter near the cytosolic side. S2 and S3 contain many aromatic residues, which face each other to interdigitate the dimer interface at the TMD (**Supplementary Fig. 7a**). Notably, a membrane-parallel short α-helix (S1a) was juxtaposed to the N-terminus of S1b, which may also contribute to maintaining the TM-helix arrangement from the outside (**Fig. 2b**). These modes of TM-helix interactions are unique to hZnT7 and have not been observed for other Zn^2+^ transporters of known structure (**Supplementary Fig. 8**).

The cytosolic domain (CTD) of hZnT7 is composed of residues 302-376 and follows the C-terminus of S6 via a short linker composed of residues Met^294^ to Leu^301^. Cryo-EM analysis revealed that the CTD comprises two α helices (α1 and α2) and four β-strands (β1a, β1b, β2, and β3) (**Fig. 1** and **Fig. 2b**), with the latter also providing a contact surface for dimerization (**Fig. 2e**). An EQ linker (Glu^323^ and Gln^324^) flanked by β1a and β1b is conserved among hZnT7 orthologues, and participates in the hydrogen bond network, along with neighboring residues of another protomer; mainchain of Glu^323^ and His^325^ from hydrogen bond to Tyr^368^ of another protomer (**Fig. 2e**). Tyr^98^ is hydrogen bonded to Gln^366^ (α2-β3 loop) and forms a π-π stack with Trp^327^ (β1b) at the TMD-CTD interface (**Fig. 2e** and **2**f). In addition, the N-terminus of S3, the C-terminus of S6, and the CTD form a hydrophobic cluster that includes Leu^106^ (S3), Leu^293^ (S6), and Trp^327^ (β1b), with hydrogen bonds between Arg^102^ and Gln^295^ further stabilizing the CTD dimer interface (**Fig. 2f**).

### Structural comparison between outward- and inward-facing forms of hZnT7

The repeated 3D classification based on cryo-EM data generated a minor class of density map at a resolution of 3.4 Å (class 2, 22%, 29,774 particles), suggesting a different conformation of hZnT7 (**Fig. 2a** and **Supplementary Fig. 3**). The cryo-EM map of this class displayed significant density for both the TMD and CTD, allowing model building for almost the entire region, including residues 22-376 (**Fig. 2b** and **Supplementary Figs. 3** and **5**). Importantly, the position and orientation of the transmembrane helices in this minor class differed significantly from those in the major class (**Fig. 2b** and **2**g). S4 and S5 are located so that the Zn^2+^ transport cavity formed by S2, S3, and S5 in the minor class is wider on the cytosolic side and narrower at the luminal side than in the major class (**Fig. 2b** and **2**g). These findings suggested that this minor class represents the inward-facing (IF) form of hZnT7 whereas the major class corresponds to its OF form. Again, the density map of the IF form showed no Zn^2+^ bound at the Zn^2+^-binding site (**Fig. 2d**).

**Fig. 5.**
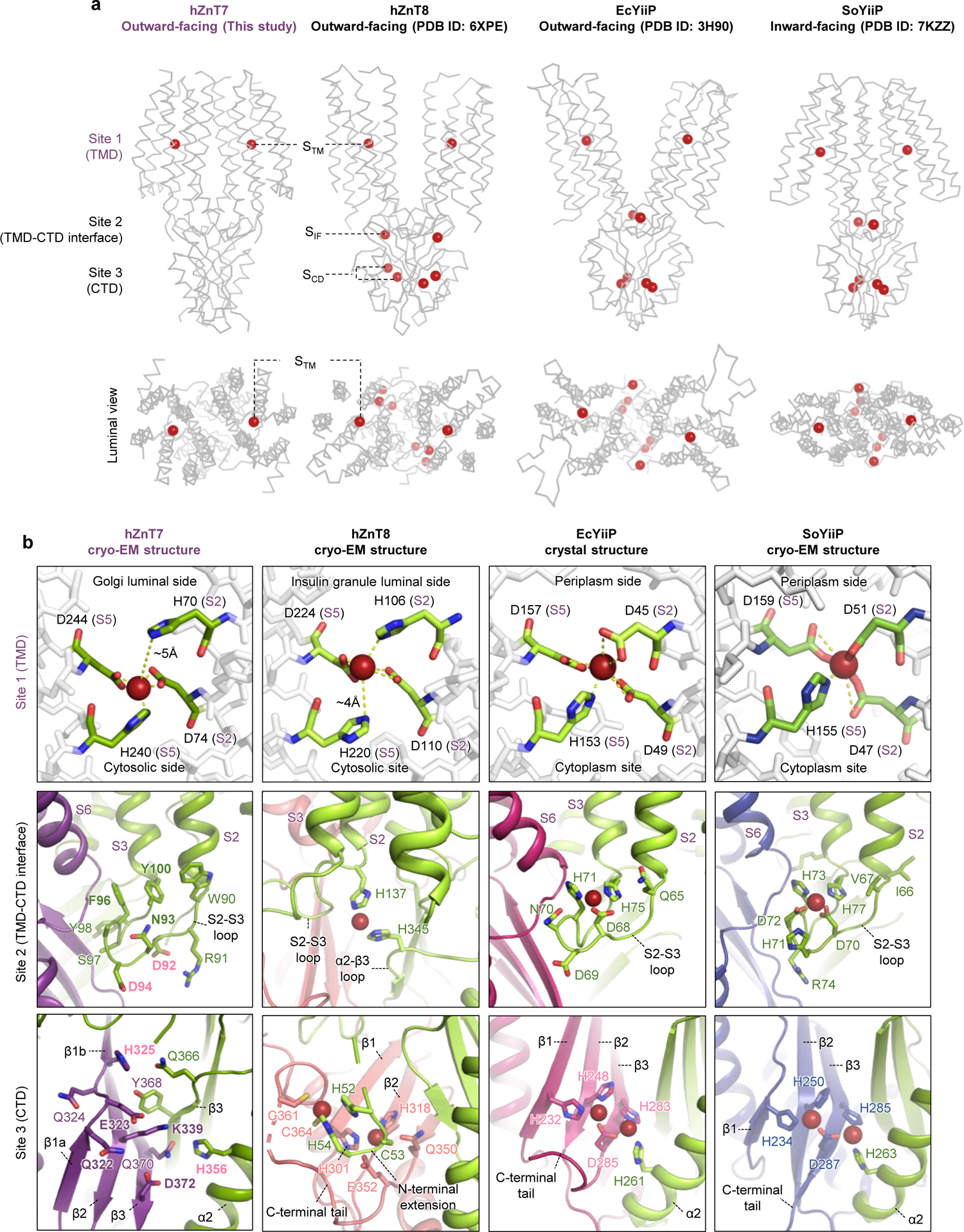
Comparison of Zn^2+^-binding sites in hZnT7 (cryo-EM structure at 2.8 Å resolution), hZnT8 (cryo-EM structure at 4.1 Å resolution), EcYiiP (crystal structure at 2.9 Å resolution), and SoYiiP (cryo-EM structure at 3.4 Å resolution). (a) Distribution of Zn^2+^-binding sites in the overall structures of hZnT7, hZnT8, EcYiiP, and SoYiiP. (b) Zinc coordination structures at sites 1, 2, and 3. Yellow dashed lines indicate Zn^2+^ coordination of the neighboring His/Asp residues in the HDHD and HDDD motifs.

Notably, the electrostatic potential surface at pH 7.5 revealed that the cytosolic gate of the Zn^2+^ transport pore near the TMD-CTD interface is negatively charged in the IF form (**Fig. 2c**), suggesting that this form can recruit Zn^2+^ from the cytosol into the cavity. By contrast, the corresponding site is positively charged in the OF form (**Fig. 2c**), consistent with this form having a closed cytosolic gate (**Fig. 2b**). S2, S3, and S5 were found to be tightly bundled at the cytosolic half of the TMD, occluding the space between the putative Zn^2+^-binding site and the cytosolic gate. By contrast, the cytosolic halves of S2, S3, and S5 in the IF form are wide open to act as a Zn^2+^-recruiting gate.

During conversion between its OF and IF forms, hZnT7 appears to undergo opposing movements of S4 and S5 at the luminal and cytosolic sides, with three leucine residues (Leu^239^, Leu^242^, and Leu^246^) in S5 apparently playing critical roles. The Leu^242^ residue in S5 seems to serve as a pivot for S5 rotation during the transition from the OF to IF forms (**Fig. 2g**). The movements of S3 and S6 are smaller on the cytosolic than on the luminal side, likely because of the tight hydrophobic interactions involving Leu^106^ (S3), Leu^293^ (S6), Tyr^98^ (S2-S3 loop) and Trp^327^ (β1b) (**Fig. 2f**), and the polar interaction between Arg^102^ at the C-terminus of S3 and Gln^295^ in the short S6-CTD loop (**Fig. 2f**). The CTDs of the OF and IF forms were almost superimposable, with a root mean square deviation (RMSD) for all Cα atoms of 0.569 Å (**Supplementary Fig. 9a**), suggesting that a conformational change in the CTD is not required for conversion between the OF and IF forms.

### Overall structure of hZnT7 in the Zn^2+^-bound state

To prepare Zn^2+^-bound hZnT7 and determine its structure, hZnT7 purified in the absence of Zn^2+^ (Zn^2+^-free buffer) was incubated with 10 μM ZnSO_4_ for 10 minutes before mixing with Fab#1. Subsequently, the hZnT7-Fab#1 complex was subjected to SEC with buffer containing 10 μM ZnSO_4_. This sample was used successfully to assess the cryo-EM structure of Zn^2+^-bound hZnT7 complexed with Fab#1 at a nominal resolution of 2.9 Å as a major class (**Fig. 3a, Supplementary Figs. 10** and **11**, and Table **1**). Another minor class (class 2, 25.4%, 48,046 particles) was generated, yielding a 3.9 Å resolution density map. Detailed comparison showed that the conformation of the minor class was highly similar to that of the major class (**Supplementary Fig. 10**). The density map of the major class displayed significant extra density near the HDHD motif comprising His^70^ (S2), Asp^74^ (S2), His^240^ (S5), and Asp^244^ (S5), indicating that Zn^2+^ binds to the TMD of hZnT7 (**Fig. 3b**). Importantly, repeated rounds of 3D classification showed that Zn^2+^-bound hZnT7 formed only an OF form, suggesting that the OF form is energetically favored in the Zn^2+^-bound state.

The overall structure of Zn^2+^-bound hZnT7 again demonstrated a “*mushroom*”-shaped dimeric architecture with tight interactions between TMDs and CTDs from two protomers (**Fig. 3a**). Calculations of electrostatic potential surface at pH 7.5 showed that the cytosolic Zn^2+^ entry site was positively charged in Zn^2+^-bound hZnT7 (**Fig. 3c**), consistent with the closed cytosolic gate observed in the OF form. Structural comparison between the Zn^2+^-bound and -unbound OF forms revealed that, upon Zn^2+^ bonding, the luminal ends of S3 and S5 slightly but significantly moved to narrow the Zn^2+^ exit gate (**Fig. 3d**). By contrast, the cytosolic ends of all TM helices barely moved due to interactions among the three bundled helices (S2, S3, and S5) on the cytosolic side (**Fig. 3d**). Zn^2+^-dependent conformational changes were negligible also for the CTD, with a RMSD for all Cα atoms in this domain of 0.212 Å (**Supplementary Fig. 9b**).

Bacterial Zn^2+^ transporters have been reported to possess a hydrophobic gate in the TMD, composed of Leu^152^ and Met^197^ for EcYiiP and of Leu^154^ and Leu^199^ for SoYiiP, to regulate Zn^2+^ entry (Fig. 1)**^18,20^**. The present study revealed that a similar leucine pair is present near the cytosolic Zn^2+^ entry gate in hZnT7. The cryo-EM structures of the OF form of hZnT7 showed that the side chains of Leu^239^ (S5) and Leu^285^ (S6) are located in close proximity (**Fig. 3e** middle and right, **Supplementary Fig. 9c**, and **Supplementary Fig. 12a** middle and right) and are engaged in tight interactions between S5 and S6 (**Fig. 2g**, and **Fig. 3d**). These findings suggest the generation of a Zn^2+^-occluded state in the OF form before transition to the Zn^2+^-releasing step. In the IF form, however, Leu^239^ (S5) is largely separated from Leu^285^ (S6) due to the movement of S5, leading to the full opening of the cytosol-facing cavity for Zn^2+^ entry (**Fig. 3e** left, **Supplementary Fig. 9c**, and **Supplementary Fig. 12a** left). Thus, the Leu^239^ (S5)-Leu^285^ (S6) pair likely acts as a switching gate that regulates Zn^2+^ entry from the cytosol depending on the OF or IF form.

### Zn^2+^ transport pathway of hZnT7

The cryo-EM structures of hZnT7 in the Zn^2+^-unbound OF and IF forms showed that a Zn^2+^/H^+^ transport pore is present in each promoter (**Fig. 4a**). Overall, this pore can be separated into two regions, the luminal and cytosolic cavities, around its halfway point, where a Zn^2+^-binding site is located (**Fig. 4a**). In the OF form, the cytosolic cavity is fully closed with a pore radius of less than 1 Å, blocking entry of the molecule/ion from the cytosolic side (**Fig. 4b**). By contrast, the luminal cavity is significantly wider, with a pore radius of 2-4 Å, allowing the access or release of molecules/ions on the Golgi luminal side. A Zn^2+^ transport pore with similar overall shape is formed in the Zn^2+^-bound state (**Fig. 4a**), but the pore radius on the luminal side is smaller than that in the Zn^2+^-unbound state of the OF form (**Fig. 4b**). Thus, Zn^2+^ binding made the luminal cavity narrower. As expected, the IF form has a wider Zn^2+^ transport pore on the cytosolic side due to the outward movement of S5 (**Fig. 4a, Fig. 2b** and **2g**), likely allowing Zn^2+^ to enter from the cytosol.

The electrostatic potential around the luminal gate of hZnT7 in the OF form, which runs downward from the luminal gate to the pore center, was found to be negative (**Fig. 4c** left and middle). Although the negatively charged luminal gate was conserved in the IF form (**Fig. 4c** right), the charge distribution in the IF and OF forms differed markedly. In the IF form, the negatively charged surface on the cytosolic cavity is exposed due to the wide opening of the cytosolic gate, consistent with the presence of multiple acidic residues near the cytosolic ends of S3 (Glu^104^) and S5 (Glu^288^). These negatively charged residues are buried inside the molecule in the Zn^2+^-bound and -unbound OF forms (**Fig. 4a**), further indicating that this region acts as a switchable Zn^2+^ recruiting gate.

### Zinc-binding site of hZnT7

The cryo-EM map of hZnT7 in the presence of Zn^2+^ demonstrated that the TMD contains a Zn^2+^-binding site (named S_TM_) at around the halfway point of the TMD (**Fig. 3a**,b and **Fig. 5a**). The S_TM_ is surrounded by the carboxyl groups of Asp^74^ (S2) and Asp^244^ (S5), and the imidazole rings of His^70^ (S2) and His^240^ (S5) (**Fig. 5b**). In the Zn^2+^-bound OF form, the Nε atom of His^70^ is distant (∼5.0 Å) from Zn^2+^ and does not directly coordinate with Zn^2+^ (**Fig. 5b**). In the Zn^2+^-unbound OF form, the side chain of Asp^74^ is turned away, further separating it from His^70^ (**Fig. 2d**) and likely leading to the facilitated Zn^2+^ release from S_TM_. In this situation, His^240^ seems to be protonated to form a hydrogen bond with the carboxyl group of Asp^74^ and stabilize the Zn^2+^ unbound conformation. By contrast, the cryo-EM structure of the Zn^2+^-unbound IF form demonstrated that, although the side chain of His^70^ is more isolated from S_TM_ (**Fig. 2d**), the other three residues, Asp^74^, His^240^, and Asp^244^, are more closely clustered, seemingly ready to accommodate Zn^2+^ entering from the cytosolic cavity.

Zn^2+^-binding sites have been observed at similar positions in the TMDs of other Zn^2+^ transporters, including bacterial YiiPs (EcYiiP and SoYiiP) and human ZnT8 (**Fig. 5b** top panels). EcYiiP and SoYiiP possess a DDHD motif that binds Zn^2+^, whereas hZnT7 and hZnT8 possess an HDHD motif at similar positions (**Fig. 5a**). Notably, the Zn^2+^ coordination structures of hZnT7 and hZnT8 differ significantly. In hZnT7, Asp^74^, His^240^ and Asp^244^ form a compact complex structure that binds Zn^2+^, while an Nε atom of His^70^ weakly coordinates with Zn^2+^ at a distance of ∼5 Å. In hZnT8, His^106^, a residue corresponding to His^70^ of hZnT7, forms a complex structure with Asp^110^ and Asp^224^ to bind Zn^2+^, whereas His^220^ is located ∼4 Å from Zn^2+^ (**Fig. 5b**). These structural findings may suggest a sequential Zn^2+^ shuttle in the interior of the S_TM_, in which His^240^ (His^220^ in hZnT8), Asp^74^ (Asp^110^ in hZnT8), and Asp^244^ (Asp^224^ in hZnT8) of hZnT7 form a triad to accept a Zn^2+^ ion from the cytosolic cavity, with a second Zn^2+^-binding triad, consisting of His^70^ (His^106^ in hZnT8), Asp^74^, and Asp^244^ being formed before releasing of Zn^2+^ on the luminal cavity. A regular tetrahedral Zn^2+^ coordination structure may transiently be formed as an intermediate between these two states, as seen in the crystal/cryo-EM structures of EcYiiP/SoYiiP (**Fig. 5b** right top).

Evaluation of the crystal structures of the Zn^2+^-bound OF form of EcYiiP (PDB ID: <u>3H90</u>) and the cryo-EM structures of the Zn^2+^-bound IF form of SoYiiP (PDB ID: <u>7KZZ</u>) and the Zn^2+^-bound OF form of human ZnT8 (PDB ID: <u>6XPE</u>) demonstrated the presence of other Zn^2+^-binding sites (**Fig. 1** and **Fig. 5a**). The second Zn^2+^-binding site of hZnT8 consists of His^137^ (S2-S3 loop) and His^345^ (α2-β3 loop) at the TMD-CTD interface (S_IF_), although this site had lower affinity for Zn^2+**<u>19</u>**^. Similarly, Zn^2+^-binding sites of EcYiiP and SoYiiP have been observed near the TMD-CTD interface (S2-S3 loop). By contrast, hZnT7 lacks a second Zn^2+^-binding site around the TMD-CTD interface, consistent with the absence of histidine residues in this region (**Fig. 5b** and **Supplementary Fig. 12b**).

Unlike hZnT7, hZnT8 and YiiP have two additional Zn^2+^ binding sites in the CTD. hZnT8 contains two cysteines near the C-terminus, and a unique HCH (His^52^-Cys^53^-His^54^) motif in a non-conserved N-terminal extension of the neighboring protomer, whereas YiiP binds two Zn^2+^ ions, S_CD1_ and S_CD2,_ in a classical trigeometry containing the (HHD)_2_ motif at the CTD dimer interface (**Fig. 5b**, bottom). The Zn^2+^-binding residues in the CTDs of YiiP and hZnT8 are replaced by other residues in hZnT7 (**Fig. 1, Fig. 5b, Supplementary Fig. 12c**, and **Supplementary Fig. 13**). Despite the absence of Zn^2+^ binding sites in its CTD, hZnT7 forms a stable homodimer in either the OF or IF form (**Supplementary Fig. 9**). Thus, Zn^2+^ ions bound to the TMD-CTD interface and the CTD seem unnecessary for the formation of functional ZnT dimers.

### Structural and functional roles of the histidine-rich loop of hZnT7

The histidine-rich loop (His-loop) of hZnT7, which is flanked by S4 and S5 on the cytosolic side, is exceptionally long (**Fig. 1**). The cryo-EM density map at 2.8 Å resolution allowed resolution of only the N-terminal and C-terminal segments of the His-loop near the ends of S4 and S5, respectively (**Fig. 2a**,b and **Fig. 3a**). Thus, the major portion of the His-loop was invisible on the cryo-EM map, suggesting that this loop is highly flexible. To understand the structural and functional roles of the His-loop, a major portion of this loop (residues His^176^ to Pro^221^) was deleted, and the resultant mutant protein (hZnT7ΔHis-loop) was overproduced in HEK293T cells and purified, as described for wild-type hZnT7, to determine its structure and perform biochemical assays (**Supplementary Fig. 14**). The cryo-EM structures of the hZnT7ΔHis-loop-Fab#1 complex in Zn^2+^-bound and -unbound states were determined at nominal resolutions of 3.4 Å (Supplementary Figs. **15**-**18** and Table **1**). Class 4 was the major class (67.4%, 127,941 particles), but displayed unusually bent TM helices and a poor density map (**Supplementary Fig. 15**). Therefore, the second major-class density map (32.6 %, 69,877 particles) was used for model building of the hZnT7ΔHis-loop.

Both Zn^2+^-bound and -unbound forms of the hZnT7ΔHis-loop were outward-facing, with similar location and orientation of the TM helices (**Fig. 6a**), indicating that removal of the His-loop had little effect on the TMD structure of hZnT7 (**Fig. 6b**-d). The CTDs of the Zn^2+^-unbound and -bound forms were also almost completely superimposable, with an RMSD for all Cα atoms in this domain of 0.204 Å (**Supplementary Fig. 19**). Collectively, although the His-loop was suggested to contribute to the adoption of the IF form of full-length hZnT7 (**Fig. 2a**), its deletion did not appreciably affect the overall structure of hZnT7.

**Fig. 6.**
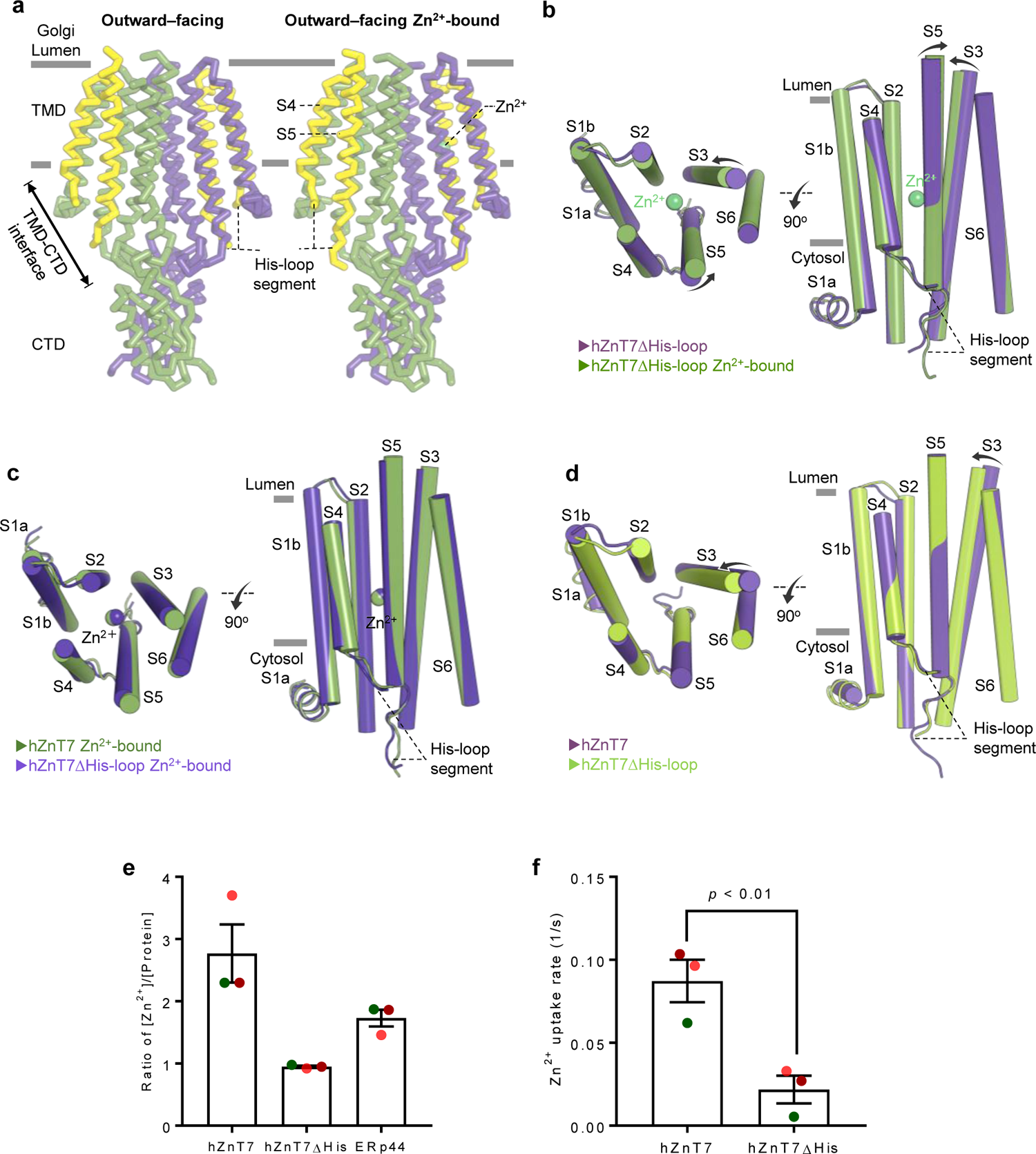
Structures and functional properties of hZnT7ΔHis-loop. (**a**) Cryo-EM structures of an outward-facing form of hZnT7ΔHis-loop in the Zn^2+^-bound and -unbound states. Transmembrane helices 4 (S4) and 5 (S5) that flank the His-loop are colored yellow. Zinc ions are indicated by green spheres. (**b**) Comparison of TMD structures in Zn^2+^-bound (purple) and -unbound (green) hZnT7ΔHis-loop. Zn^2+^ is shown as a green sphere. (**c**) Comparison of TMD structures of the outward-facing forms of Zn^2+^-bound hZnT7ΔHis-loop (green) and hZnT7 WT (blue). (**d**) Comparison of TMD structures of the outward-facing forms of Zn^2+^-unbound hZnT7ΔHis-loop (lime) and full-length hZnT7 (purple). (**e**) Zinc-binding stoichiometry of purified hZnT7, hZnT7ΔHis-loop, and ERp44 (control) determined by a spectroscopic assay using Zincon. Data are from three independent experiments. Error-bars indicate mean ± SEM. (**f**) Zinc uptake rate of hZnT7 WT and hZnT7ΔHis-loop determined by incubating ZnT7-embedded proteoliposomes containing Fluozin3 in buffer containing 100 μM ZnSO_4_. Data are from three experiments. Error-bars indicate mean ± SEM.

To predict the location and structure of the His-loop that was invisible in cryo-EM map, a model of the entire part of the hZnT7 homodimer was generated using AlphaFold2**^39^**. Only an OF form structure was generated by this program, which showed the similar TM helix arrangement and CTD fold to those in the cryo-EM structure of Zn^2+^-unbound hZnT7 (**Supplementary Fig. 20**). Histidine residues were predicted to localize near the C-terminus of S4 and to be distributed throughout the His-loop (**Supplementary Fig. 20a**). Importantly, the histidine residues near the C-terminus of S4 were likely located near the cytosolic Zn^2+^-entry gate. We surmise that the His-loop with such a histidine distribution pattern is likely essential for efficient Zn^2+^ binding and transport by hZnT7.

To test this possibility, the Zn^2+^-binding abilities of hZnT7ΔHis-loop and wild-type hZnT7 (hZnT7 WT) were compared. The Zn^2+^-binding stoichiometry of hZnT7ΔHis-loop, hZnT7 WT, and ERp44 as a control were measured by quantifying bound Zn^2+^ with the zinc probe Zincon. ERp44 bound 1.7 ± 0.1 molar equivalents of Zn^2+^ (**Fig. 6e**), in agreement with our previous observation that ERp44 forms a homodimer with three high-affinity Zn^2+^-binding sites^40^. Similarly, the Zn^2+^-to-protein molar ratios were determined to be 2.8 ± 0.5 for hZnT WT and 1.0 ± 0.0 for the hZnT7ΔHis-loop (**Fig. 6e**). These findings indicate that hZnT7 binds two extra Zn^2+^ ions via its His-loop in addition to the one in the TMD (**Fig. 3a** and **Fig. 6a**). The role of the His-loop in Zn^2+^ transport was assessed by performing a Zn^2+^ transport assay using proteoliposomes containing a Fluozin-3 probe inside. Deletion of the His-loop was found to significantly impair the Zn^2+^ transport activity of ZnT7 (**Fig. 6f**). These results suggest that the His-loop of hZnT7 is essential for recruiting Zn^2+^ ions from the cytosol and transporting them to the Golgi lumen.

### Structure comparisons of hZnT7 with other Zn^2+^/H^+^ transporters

Comparisons of hZnT7, hZnT8, SoYiiP, and EcYiiP revealed marked differences in their overall structures. hZnT7 dimers in the OF form formed a “*mushroom*”-shaped dimeric structure with tight interactions between S2 and S3 from two different protomers (Supplementary Figs. **7** and **8**). By contrast, hZnT8 and EcYiiP dimers formed a “*V*” shaped dimeric structure with much less tight interactions between the TMDs from the two protomers (Fig. 5a)^14, 19^. In this context, although hZnT7 formed homogeneous dimers only with both protomers in the IF or OF form^14–18^, hZnT8 could form a heterogeneous structure, with one protomer in the IF form and another in the OF form^19^.

**Fig. 7.**
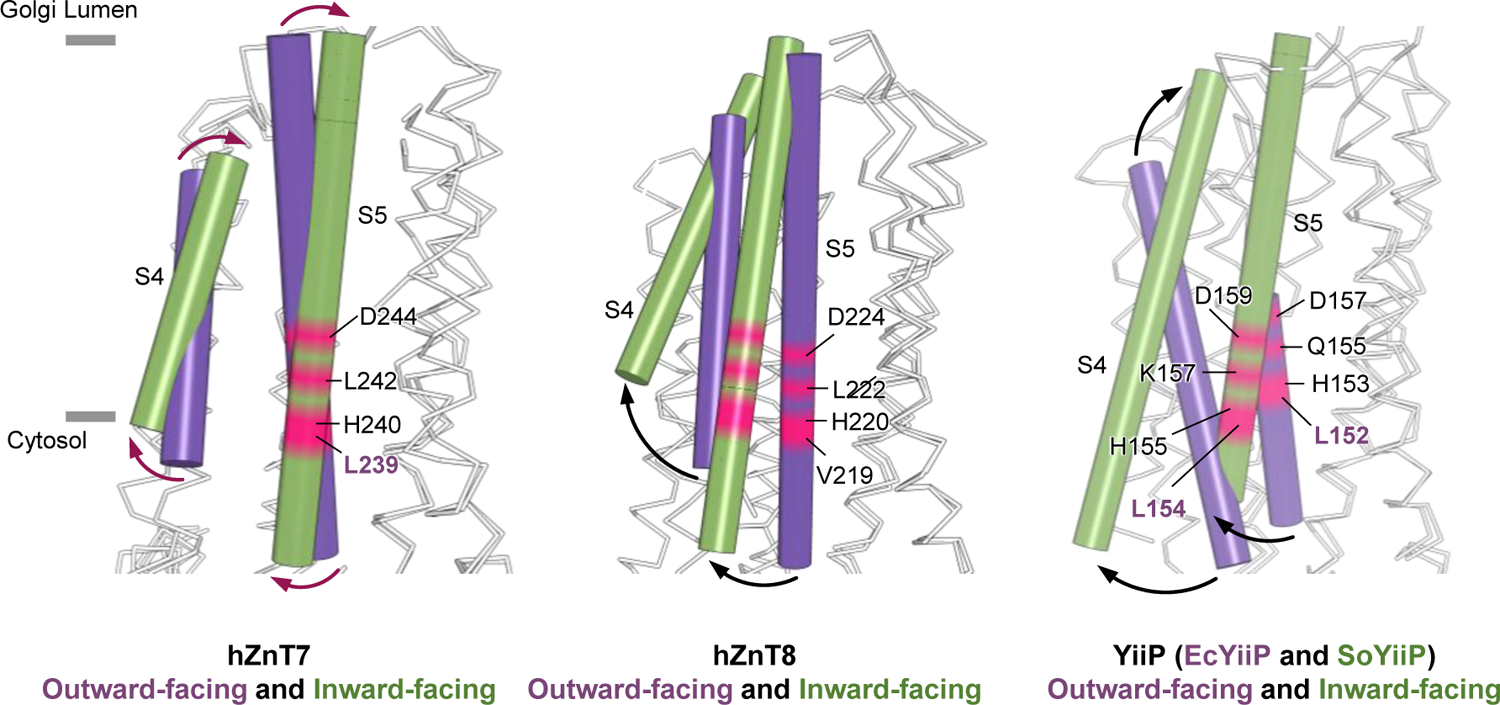
Comparison of TMD structures of hZnT7, hZnT8, and bacterial YiiP. Comparisons of TMD structures of the OF and IF forms of hZnT7 (left), the OF (PDB ID: <u>6XPE</u>) and IF (PDB ID: <u>6XPF</u>) forms of hZnT8 (middle), and the OF form of EcYiiP (PDB ID: <u>3H90</u>) with the IF form of SoYiiP (PDB ID: <u>7KZZ</u>) (right). S4 and S5 were found to move to different degrees during the conversion of hZnT7 (left), hZnT8 (middle), and YiiP8 (right) from the OF to the IF form. For clarity, S4 and S5 are represented by cylinders.

**Fig. 8.**
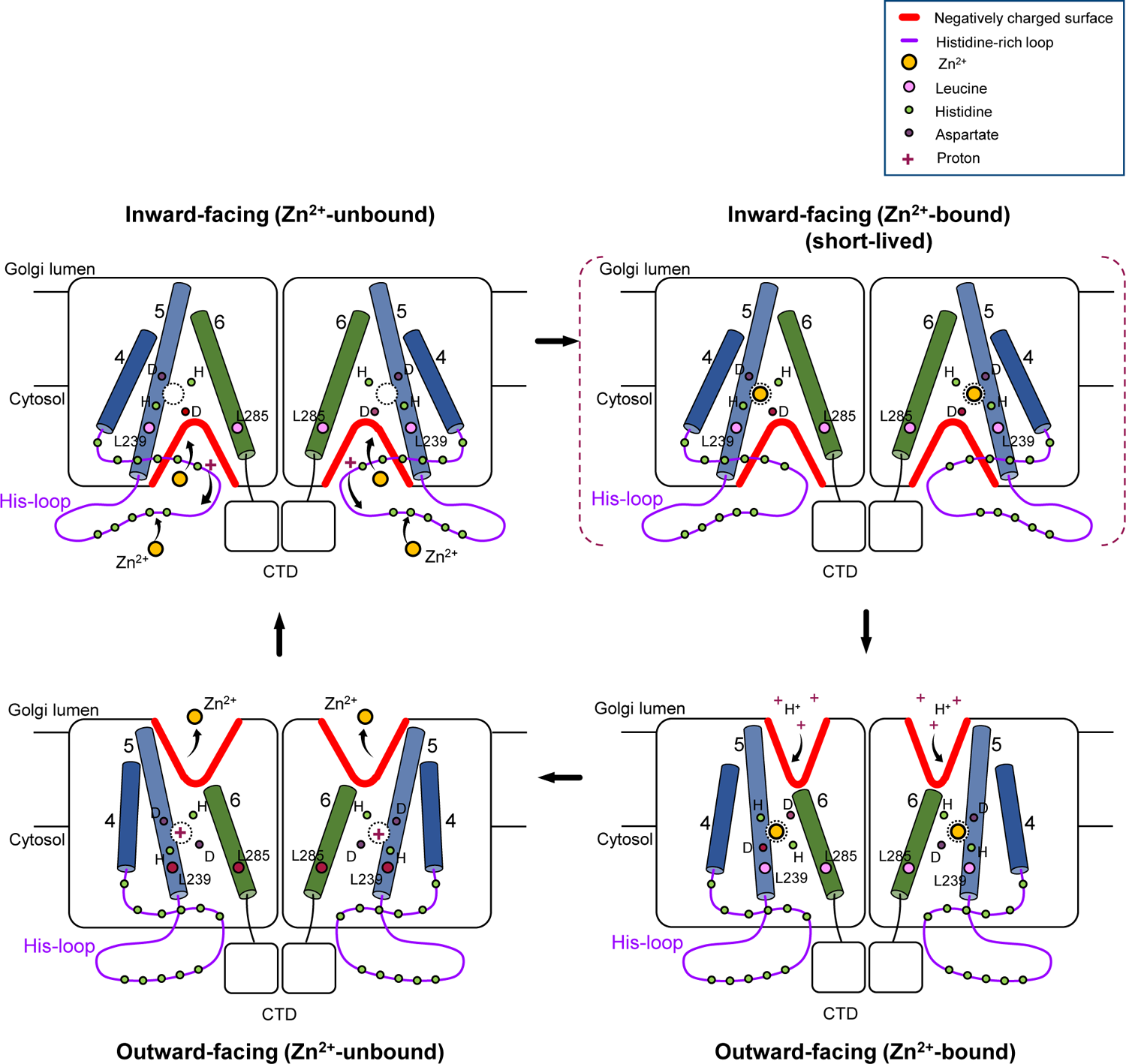
Proposed model of Zn^2+^ transport mediated by a Golgi-resident human Zn^2+^/H^+^ antiporter hZnT7. During its Zn^2+^ transport cycle, hZnT7 undergoes TM-helix rearrangement, resulting in a negatively charged cytosolic cavity for Zn^2+^ entry in the IF form and a negatively charged luminal cavity for H^+^ incorporation and Zn^2+^ release in the OF form. The exceptionally long His-loop may facilitate Zn^2+^ recruitment from the cytosol and subsequent Zn^2+^ shuttling to the cytosolic cavity.

Superimposition of the cytosolic domains of the OF and IF forms showed that S4 and S5 of hZnT7, hZnT8, and EcYiiP move differently during the conversion between these two forms. S4 and S5 of hZnT7 rotate around their middle regions, whereas S4 and S5 of hZnT8 swing using their luminal ends as pivot points (**Fig. 7**). Consequently, the cytosolic ends of S4 and S5 move to a greater extent in hZnT8 than in hZnT7, generating a wider cytosolic cavity in the IF form of hZnT8. Although only the structures of the OF form of EcYiiP and the IF form of SoYiiP have been solved, superimposition of their TMDs suggests that S4 and S5 of the bacterial YiiP rotate to an even greater extent than hZnT7, thereby generating a larger cytosolic cavity (**Fig. 7**). In conclusion, hZnT7 undergoes smaller TM-helix movements during the transition between the OF and IF forms than do other zinc transporters of known structures.

In this context, a short membrane-parallel α-helix at the N-terminus of S1 (S1a) of hZnT7 was found to be located on the cytosolic side, with small movement between the OF and IF forms (**Fig. 2g**, left). Neither hZnT8 nor YiiP possesses such a membrane-parallel α-helix element on the cytosolic side (**Supplementary Fig. 8**). The S1a segment characteristic of hZnT7 may limit the movement of the cytosolic regions of the TM helices during the transition from the IF to OF form.

### Mechanism of hZnT7-mediated Zn^2+^ transport into the Golgi lumen

Altogether, the cryo-EM and biochemical analyses in the present study provide a likely scenario for hZnT7-mediated Zn^2+^ transport from the cytosol to the Golgi lumen (**Fig. 8**). The Zn^2+^-unbound IF form of hZnT7 possesses a negatively charged cytosolic cavity at the TMD-CTD interface, where the exceptionally long and flexible His-loop is predicted to localize. These structural features seem advantageous for the efficient recruitment and shuttling of Zn^2+^ to the negatively charged cytosolic Zn^2+^ entry gate. The hydrophobic gates in EcYiiP (Leu^152^ and Met^197^) and SoYiiP (Leu^154^ and Leu^199^) had been found to regulate Zn^2+^ entry**^18,20^**. Similarly, the present study revealed that a leucine-pair gate (Leu^239^ and Leu^285^) is present near the cytosolic Zn^2+^ entry gate in hZnT7. The hydrophobic gate was fully open in the Zn^2+^-unbound IF form, with Zn^2+^ binding triggering the closure of the leucine-pair gate to generate a Zn^2+^-occluded state before transition to the Zn^2+^-releasing state (i.e. Zn^2+^-bound OF form) via TM-helix rearrangements.

The Zn^2+^-bound OF form has structural features suitable for Zn^2+^-release to the Golgi lumen. The luminal cavity of the Zn^2+^ transport pore is open, whereas the cytosolic cavity is fully closed to prevent further Zn^2+^ entry from the cytosol. Presumably, H^+^ from the weakly acidic Golgi lumen protonates residues of the HDHD motif, triggering the release of bound Zn^2+^. The negatively charged surface of the luminal cavity likely facilitates Zn^2+^ transfer to the luminal gate. Protonations at the HDHD motif may also induce the subsequent conversion from the OF to the IF form, releasing H^+^ into the cytosol. This mechanism of action nicely explains the physiological function of hZnT7 as a Zn^2+^/H^+^ antiporter in the Golgi membrane. ZnT7 is likely to collaborate with other ZnT and ZiP family members for the maintenance of Zn^2+^ homeostasis in the early secretory pathway. Detailed mechanisms of Zn^2+^ homeostasis resulting from the cooperation of multiple Zn^2+^ transporters remain to be elucidated, in light of the physiological roles of Zn^2+^ in the folding and maturation of Zn^2+^-binding proteins, cell signaling and regulation, and many other life activities.

## Methods

### Cloning and protein expression

A cDNA encoding human ZnT7 (hZnT7) was synthesized with codon-optimization for expression in human cells. The hZnT7 gene with an N-terminal PA-tag sequence (GVAMPGAEDDVV) was cloned into a PiggyBac Cumate Switch Inducible Vector (SBI). For large-scale expression of hZnT7, stable PiggyBac/hZnT7-expressing HEK293T cells were cultured in DMEM (High glucose; 4.5 g/L D-(+)-glucose with L-glutamine, phenol red, and sodium pyruvate; Nacalai Tesque), supplemented with 4% fetal bovine serum (FBS; Biosera), 1% penicillin-streptomycin mixed solution (PS; Nacalai Tesque), and 4 μg ml^-1^ puromycin (InvivoGen), and grown at 37 °C in an atmosphere containing 5% CO_2_. After 3 days, the cells were passaged into fresh DMEM (high glucose) supplemented with 4% FBS, and 1% PS. After growth for 24 h, expression was induced by addition of Cumate (Wako) and phorbol 12-myristate 13-acetate (Wako) to final concentrations of 10X (300 μg ml^-1^) and 1X (50 nM), respectively. Cells were continuously cultured at 37 °C for 3 h in 5% CO_2_ before being incubated at 30 °C in 5% CO_2_. After 48 h, the cells were harvested, washed with cold-1X phosphate buffered saline (PBS), and stored at −80°C.

### Protein purification

Proteins were purified and treated at 4°C. Cells were washed with ice-cold-1X PBS buffer and collected by centrifugation. Cells from 2 liters of medium (wet-weight, ∼8 grams) were lysed by resuspension in buffer A containing 20 mM Tris-HCl, pH 7.5, 150 mM NaCl, and 10% glycerol, supplemented with 2 mM DTT (Nacalai Tesque), 0.05 mg ml^-1^ DNase I (Wako), 5 mM MgCl_2_ (Wako), and 1/100 protease inhibitor cocktail (Nacalai Tesque). The cells were sonicated for 3 min in an ice-water cup, and the cell membrane fraction was collected by ultracentrifugation at 200,000 ×g (micro Ultracentrifuge CS100FNX, Hitachi) for 1 h at 4 °C. The pellet was dissolved in a solution containing 1% (v/v) n-dodecyl-β-D-maltoside (DDM; Nacalai Tesque), 0.5% (v/v) cholesteryl hemisuccinate (CHS; Sigma), 20 mM Tris-HCl, pH 7.5, 150 mM NaCl, 10% glycerol, and 2 mM DTT, and incubated for 2 h at 4 °C with constant stirring. The supernatant containing PA-tagged hZnT7 was collected by centrifugation at 15,000 ×g for 30 min at 4 °C and incubated with 12 ml (net weight, 6 ml) Anti-PA-tag Antibody Beads (Wako) overnight at 4 °C with gentle rotation. The resin was collected in a glass column (BioRad) and washed with 15 CV of Buffer B [buffer A supplemented with 0.02% GDN (w/v) (Anatrace), and 2 mM DTT (w/v)]. PA-tagged hZnT7 was eluted with Buffer C [Buffer B supplemented with 0.2 mg ml^-1^ PA peptide (PH Japan Co., Ltd)]. The protein was concentrated using a 50-kDa cutoff Amicon filter (Sigma-Aldrich) and purified on a Superose 6 Increase 10/30 gel filtration column (GE Healthcare), pre-equilibrated with buffer A supplemented with 2 mM DTT (v/v), and 0.02% GDN (w/v). The eluted fractions were concentrated using a 100-kDa cutoff Amicon filter to ∼10 mg ml^-1^ for negative staining EM and cryo-EM measurements.

### Preparation and screening of monoclonal antibody Fab fragments to hZnT7

All animal experiments conformed to the guidelines of the Guide for the Care and Use of Laboratory Animals of Japan and were approved by the Kyoto University Animal Experimentation Committee. Mouse monoclonal antibodies against hZnT7 were generated essentially as described^21^. Briefly, a proteoliposome antigen was prepared by reconstituting purified hZnT7 at high density into phospholipid vesicles consisting of a 10:1 mixture of chicken egg yolk phosphatidylcholine (egg PC; Avanti Polar Lipids) and adjuvant lipid A (Sigma-Aldrich) to facilitate the immune response. MRL/lpr mice were injected three times at 2-week intervals with the proteoliposome antigen. Antibody-producing hybridoma cell lines were generated using a conventional fusion protocol. Biotinylated proteoliposomes were prepared by reconstituting hZnT7 with a mixture of egg PC and 1,2-dipal-mitoyl-sn-glycero-3-phosphoethanolamine-N-(cap biotinyl) (16:0 biotinyl Cap-PE; Avanti), and used as binding targets for conformation-specific antibody selection. The targets were immobilized onto streptavidin-coated microplates (Nunc). Hybridoma clones producing antibodies recognizing conformational epitopes of hZnT7 were selected by enzyme-linked immunosorbent assay on immobilized biotinylated proteoliposomes (liposome ELISA), allowing positive selection of the antibodies that recognized the native conformation of hZnT7. Clones were additionally screened for reduced antibody binding to SDS-denatured hZnT7, resulting in negative selection of linear epitope-recognizing antibodies. The formation of stable complexes between hZnT7 and each antibody clone was determined by fluorescence-detection SEC. Five monoclonal antibodies were found to specifically bind to and stabilize conformational epitopes of hZnT7. The sequences of the Fabs were determined by standard 5’-RACE using total RNA isolated from hybridoma cells.

### Preparation of hZnT7-Fab complexes

Purified hZnT7 was mixed at a molar ratio of 1:3 with each of the five selected Fab fragments (Fab#1, YN7114-08; Fab#2, YN7117-01; Fab#3, YN7114-03; Fab#4, YN-7148-12; and Fab#5, YN7179-03). Following incubation for 30 min at 4 °C with gentle rotation, the mixtures were centrifuged at 15,000 r.p.m. for 10 min at 4 °C and injected into Superdex 200 Increase 10/300 GL size-exclusion columns (GE Healthcare). Fractions containing both hZnT7 and Fab were collected and concentrated with a 100-kDa cutoff Amicon filter to around 10 mg ml^-1^ for cryo-EM measurements.

To prepare Zn^2+^-bound hZnT7, hZnT7 purified in Zn^2+^-free buffer was mixed with 10 μM Zn^2+^ and incubated on ice for 10 min. This mixture was incubated with a 3-molar excess of Fab#1 for 0.5 h at 4°C with gentle rotation, centrifuged at high speed to remove precipitates, and injected onto an SEC column, as described above, with the SEC buffer containing 10 μM Zn^2+^. Fractions containing both hZnT7 and Fab#1 were pooled and concentrated using a 100-kDa cutoff Amicon filter to ∼8 mg ml^-1^ for cryo-EM measurements.

### Preparation of hZnT7ΔHis-loop sample

To prepare the hZnT7ΔHis-loop, the DNA segment encoding residues His^176^ to Pro^221^ of hZnT7 was deleted by QuickMutagensis using the primers 5’-CCACAGCAGCCTGAAAGAGACAACC-3’ (forward) and 5’-TTCAGGCTGCTGTGGCCGTGTCCAGA-3’ (reverse), both synthesized by Eurofins. The medium was changed to fresh serum-free DMEM containing 1% PS, and HEK293T cells were transfected with pcDNA3.1/PA-hZnT7ΔHis-loop plasmids using PEI reagent (Sigma-Aldrich) at a ratio of 50 μg DNA to 1.5 ml PEI reagent (1 mg ml^-1^). Cells were continuously cultured for 24 h before 10 mM sodium butyrate (Wako) was added. After 54 h of culturing at 30 °C and 5% CO_2_, the cells were harvested. hZnT7ΔHis-loop protein was purified as described for wild-type hZnT7.

### Negative stain EM

The hZNT7-Fab complexes were assessed by negative-stain EM, which provided 2D reconstruction images useful for Fab selection. Grids for negative-stain EM were prepared according to the standard protocol. Specifically, 5 μl of purified hZnT7-Fab complex was applied to glow-discharged carbon-coated grids (ELC-C10), with the excess solution removed with filter paper. The grid was immediately stained with 1 % (w/v) uranyl acetate solution. The negatively stained EM grids were imaged on a JEOL JEM-2010F electron microscope (JEOL) operated at 200 kV. Particles were extracted by phase-flipping, followed by reference-free 2D class averaging using RELION-3.1 version**^22^**.

### EM data acquisition

To prepare cryo-EM grids, 3 μl of hZnT7-fab complex samples were applied to glow-discharged QuantiFoil R1.2/1.3 Cu/Rh 300 mesh grids (for hZnT7) or Cu 300 mesh grids (for hZnT7ΔHis-loop). The grids were blotted and immersed in liquid ethane using Vitrobot Mark IV systems (FEI/Thermo fisher) operated at 4 °C and 100% humidity. The blotting parameters for complexes of hZnT7 with Fab#1 or Fab#3 and for Zn bound hZnt7 were set at a wait time of 3 sec, a blot time of 3.5-4 sec, and a blot force of 10; and the blotting parameters for hZnT7-ΔHis-loop with or without Zn were set at a wait time of 3 sec, a blot time of 3 sec, and a blotting force of −5.

The grids were initially imaged using a Talos Arctica TEM (Thermo Fisher Scientific) equipped with a Gatan K2 summit. Subsequently, movies of the hZnT7 with Fab#1 were collected on a Titan Krios G3i TEM (Thermo Fisher Scientific) operated at 300 kV, equipped with a Gatan Quantum-LS Energy Filter (GIF) and a Gatan K3 BioQuantum direct electron detector. Movies of the hZnT7ΔHis-loop-Fab complex were collected on a CRYO ARM™ 300II (JEOL) operated at 300 kV and equipped with a JEOL in-column Omega energy filter and a Gatan K3 BioQuantum detector. Data were automatically collected using SerialEM software^23^. The data collection parameters are summarized in Table **1**.

### EM data processing

All movie stacks were subjected to motion correction using the motion correction program implemented in RELION 3.1^22^, with the contrast transfer function (CTF) parameters estimated with CTFFIND4^24^. For the dataset of hZnT7 in Zn^2+^-free buffer (Dataset 1), particles were initially picked from 1,016 movies using Laplacian-of-Gaussian-(LoG) based auto-picking in RELION3.1 and selected from 2D classification as 2D references, which were used for template-based auto-picking from 4,275 movies. 2D class averages representing hZnT7-Fab#1 complexes were separately selected from top views (i.e. from Fabs or detergent micelles) and side views, with further 2D classification performed for each view. The representative 2D classifications of hZnT7-Fab in side (295,855 particles) and top (303,643 particles) views were combined and subjected to two rounds of 3D classifications in C1 symmetry (4 to 6 classes) using the initial model generated by RELION 3.1. The best particles (145,096 particles) were re-extracted at a pixel size of 1.19889 Å and subjected to 3D refinement in C2 symmetry, CTF refinement, and Bayesian polishing. After one round of 3D classification without alignment using an overall mask, selected particles (139,367 particles) were processed with additional CTF refinement and 3D refinement, which yielded an EM map with a resolution of 2.7 Å. To improve the density for S4 and S5, focused 3D classification without alignment was performed using a mask covering the TM domain (fourth 3D classification). After removal of duplicates, particles belonging to the major class (class 2, outward form) were subjected to 3D refinement followed by non-uniform refinement in cryoSPARC. Although the resulting 3D reconstruction generated an EM map at 2.7 Å resolution, the EM densities of S1 and S4 were not clear. Subsequently, 25,077 particles were further selected from another focused 3D classification using a mask covering the S1 and S4 regions and processed with non-uniform refinement, which generated an EM map at 2.8 Å resolution. In addition, particles corresponding to a minor class in the fourth classification, which represented different conformations (class 1, inward form), were refined at 3.4 Å resolution with 3D refinement in RELION 3.1 (**Supplementary Fig. 3**).

For the dataset of hZnT7 in Zn^2+^-containing buffer (Dataset 2), particles were picked from 4,419 movies by auto-picking in RELION 3.1 using the 2D references from dataset 1. A total of 595,890 particles were selected from two rounds of 2D classification and were applied to the two rounds of 3D classification in C1 symmetry using the initial model generated by RELION as reference. The best particles (112,859 particles) were re-extracted at a pixel size of 1.08937 Å, and subjected to 3D refinement, CTF refinement, and Bayesian polishing, followed by non-uniform refinement in cryoSPARC. To improve the resolution of S1 and S4, particles were further selected from focused 3D classifications with a mask focusing only on S1 and S4, which yielded an EM map at 2.9 Å resolution. An EM map reconstructed from particles classified into a minor class (48,046 particles) at 3.9 Å resolution was similar to that of the major class (**Supplementary Fig. 10**).

For the dataset of hZnT7ΔHis-loop in Zn^2+^-free buffer (Dataset 3), particles were picked from 1,213 movies using LoG-based auto-picking in RELION 3.1 to create 2D references for template-based picking from the full dataset of 4,850 movies. After two rounds of 2D classification, 325,036 particles were selected and subjected to two rounds of 3D classification using the 3D initial model generated by RELION 3.1. Particles classified into the best class (188,918 particles) were re-extracted at a pixel size of 1.182 Å and subjected to 3D refinement in C2 symmetry, CTF refinement, and Bayesian polishing. Subsequent focused non-aligned 3D classification in C2 symmetry using an encompassing mask of the TM domain improved the local resolution of the latter. The best particles with clear density for the TM domain (Class 2, 60,977 particles) were refined at 3.4 Å resolution with non-uniform-refinement in cryoSPARC (**Supplementary Fig. 15**).

For the dataset of the hZnT7ΔHis-loop in Zn^2+^-containing buffer (Dataset 4), particles were picked from 500 movies by using LoG-based auto-picking in RELION3.1 to create 2D references for template-based picking from the full dataset of 6,150 movies. After two rounds of 2D classification, a total of 424,349 particles were selected and subjected to two rounds of 3D classification. Particles classified into the best class (142,746 particles) were re-extracted at a pixel size of 1.182 Å and subjected to 3D refinement in C2 symmetry, CTF refinement, and Bayesian polishing. Subsequent focused non-aligned 3D classification in C2 symmetry using an encompassing mask of the TM domain improved the local resolution of the latter. The best particles (142,624 particles) were refined at 3.4 Å resolution with non-uniform refinement in cryoSPARC (**Supplementary Fig. 16**).

Global resolution was estimated in RELION and cryoSPARC with a Fourier shell correlation (FCS) of 0.143. The EM maps were sharpened and locally filtered based on local resolution with cryoSPARC^25^ or sharpened with Auto-sharpen in PHENIX^26^.

### Model building, refinement, and validation

An initial model of the Zn^2+^-free hZnT7-Fab#1 complex was automatically built with Buccaneer in CCP-EM^27^. Further manual model building was performed with Coot^28, 29^. TM helices at low-resolution were modeled based on a homology model created using the SWISS model webserver. The model was refined against the locally filtered map using phenix.real_space_refine in Phenix^30^. The final model included hZnT7 residues 22-376, Fab light chain residues 1-218, and Fab heavy chain residues 1-234. The histidine-rich loop (residues 163 to 224) was missing from the EM map. hZnT7 residues 135-140 and 258-264 were removed from the model because of poor density maps. The structures of the IF form of hZnT7 and the OF form of hZnT7ΔHis-loop were initially modeled using the Zn^2+^-unbound OF structure, followed by further manual model building with Coot. The models were refined against the locally filtered maps or against phenix-auto-sharpened maps (for IF structure) using phenix.real_space_refine.

The final models were validated using MolProbity^31^. All the figures were prepared in Pymol^32^, UCSF Chimera^33^, and UCSF ChimeraX^34^. Pore radii were calculated using the HOLE program^35^ and analyzed in Pymol^32^ and VMD^36^. Amino acid sequence alignments were analyzed by the MAFFT version 7 online service with defaults^37^. Electrostatic potential maps were calculated with APBS-PDB2PQR^38^. An entire hZnT7 structure with the His-loop was predicted by AlphaFold2^39^.

### Zn^2+^-binding assay

hZnT7 and hZnT7ΔHis-loop were purified in Zn^2+^-free buffer as described above, and ERp44 was purified as described previously^40^. For Zn^2+^-binding assay, 10-20 μM protein samples were mixed with 100 μM ZnCl_2_ in Buffer A (50 mM HEPES, pH 8.0, 300 mM NaCl, 100 mM sucrose) for 10 min on ice. The hZnT7 and hZnT7ΔHis-loop samples were maintained in 0.02% GDN. Samples were applied to PD-10 columns (GE Healthcare) to remove residual Zn^2+^. Eluted fractions were collected and concentrated using 10-kDa cutoff Amicon filters. Protein concentrations were measured using BCA assay kits (Wako). A 50 μl aliquot of each sample was mixed with 80 μl denaturation buffer (50 mM HEPES, pH 8.0, 6 M guanidine hydrochloride, 10% SDS) at room temperature for 1 h, followed by incubation with 30 μM Zincon for 10 min at room temperature^41^. The absorbance of the Zn^2+^-Zincon complex in the visible region was measured using a Spectrophotometer U-3900 (Hitachi). Data were collected from three independent experiments. Purified ERp44 was used as a positive control. Zincon monosodium salt (Sigma-Aldrich) was prepared immediately before each assay. A stock solution of Zincon was prepared in MilliQ-filtered water, diluted, and the concentration was determined based on the absorbance at 488 nm using ε_488_ = 26 900 M^−1^·cm^−141^.

### Zn^2+^ transport assay with proteoliposomes

The Zn^2+^ transport activity of hZnT7 was measured using FluoZin-3 (Invitrogen), a Zn^2+^-sensitive fluorophore. To avoid bleaching of the fluorophore, the sample was shielded from direct light throughout the experiments. Briefly, 1 mg of purified hZnT7 or hZnT7ΔHis-loop was added to 5 mg egg yolk phosphatidylcholine (egg PC, Avanti Polar Lipids) in 1 mL of PBS containing 0.8% sodium cholate (Sigma). The solution was incubated with 100 mg of wet fresh Bio-Beads SM-2 (Sigma) overnight at 4 °C, with gentle rotation. The beads were removed, and the supernatant was centrifuged at 200,000 xg for 20 min at 4 °C. Each pellet was resuspended in 100 μl Buffer IN (20 mM HEPES, pH 6.5, 150 mM KCl), and mixed with 50 μM Fluozin-3 (Sigma). The mixtures were sonicated for 30 sec, frozen in liquid nitrogen, and thawed on ice. This procedure was repeated twice. The mixtures were again sonicated for 30 sec and applied to a PD-10 column. The eluted fractions were centrifuged at 200,000 xg for 20 min at 4 °C. The pellets were resuspended in 100 μl Buffer IN and kept on ice prior to the assay. As a negative control, empty liposomes were prepared using the same protocol. The proteoliposomes and liposomes thus prepared were mixed with 100 μM ZnSO_4_ in a cuvette containing 500 μl buffer OUT (20 mM HEPES, pH 7.5, 150 mM KCl). Fluorescence spectra were recorded at an excitation wavelength of 490 nm using a Fluorescence Spectrophotometer F-7000 (Hitachi). To normalize the time-dependent fluorescence, a solution of 1% OG detergent in 500 μl buffer OUT was added to each proteoliposome or liposome preparation for 1 h on ice in the dark to determine the maximal fluorescence of each individual preparation (FP_max_). Background signals from empty liposomes were used to normalize the FL_max_. Zn^2+^ transport activity was calculated as FP/FP_max_ - FL/FL_max_ over time, with initial transport rates (ΔF s^−1^) calculated by linear regression of the transport data^42^.

## Acknowledgments

We thank T. Yokoyama, K. Nanatani, J. Inoue, S. Koshiba, and M. Yamamoto for management of the cryo-EM facility at Tohoku University medical megabank. This work was supported by funding from AMED-CREST (21gm1410006h0001) to K.I., JSPS KAKENHI to K.I. (18H03978, 21H04758, and 21H05247), Canon Medical Systems Corporation to K.K. and K.I., and the Basis for Supporting Innovative Drug Discovery and Life Science Research (BINDS) from the Japan Agency for Medical Research and Development (AMED) under grant numbers JP19am0101115 (support number: 1025) and JP21am0101079 (support no. 2343).

## Author contributions

B.B.H. performed almost all experiments, structure modeling, and structure refinement. A.T. and M.K. acquired cryo-EM images. S.W. performed image processing, structure modeling and refinement, and assisted in sample preparation. N.N., K.L., T.U. and S.I. prepared the monoclonal antibody Fab fragment recognizing hZnT7. M.U. prepared a stable expression cell line for hZnT7. B.B.H. and K.I. prepared the figures and wrote the manuscript. All authors discussed the results, critically read the manuscript, and approved the manuscript for submission. K.I. supervised this work.

## Data and materials availability

The atomic coordinates of human ZnT7 have been deposited in the Protein Data Bank under accession codes <u>7X1D</u> (Zn^2+^-unbound OF form), <u>7X1P</u> (Zn^2+^-unbound IF form), <u>7X1S</u> (Zn^2+^-bound OF form), <u>7X1E</u> (ZnT7ΔHis-loop in Zn^2+^-unbound OF form), and <u>7X1F</u> (ZnT7ΔHis-loop in Zn^2+^-bound OF form). Cryo-EM density maps of human ZnT7 have been deposited in the Electron Microscopy Data Bank under accession codes <u>32937</u> (Zn^2+^-unbound OF form), <u>32945</u> (Zn^2+^-unbound IF form), <u>32948</u> (Zn^2+^-bound OF form), <u>32938</u> (ZnT7ΔHis-loop in Zn^2+^-unbound OF form), and <u>32939</u> (ZnT7ΔHis-loop in Zn^2+^-bound OF form).

## Supplementary Figures

**Supplementary Fig. 1.**
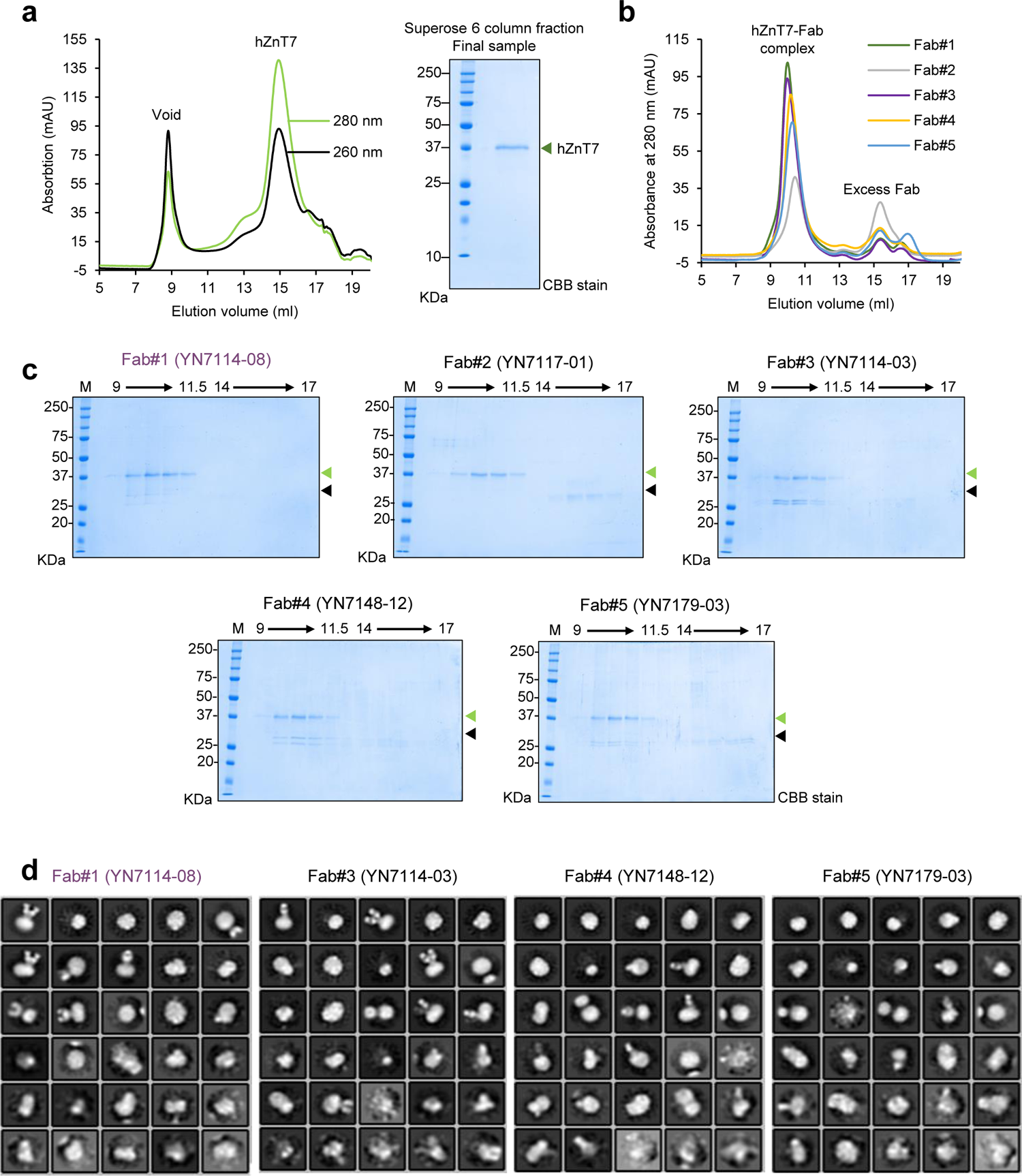
hZnT7 purification and hZnT7-Fab complex sample preparation for Cryo-EM analysis. **a**, Representative chromatogram from Superose 6 Increase gel filtration of hZnT7 (left) and a Coomassie-stained SDS gel of the final hZnT7 sample (right). The final sample consisted of gel filtration chromatography fractions at elution volumes of 13.5 to 16.5 ml. **b**, Representative chromatogram from Superdex 200 gel filtration of hZnT7 in complex with Fab fragments (#1 to #5) made from selected monoclonal antibodies. **c**, Coomassie-stained SDS gels of the hZnT7 (green triangle)-Fab (black triangle) complex separated by gel filtration chromatography shown in panel b. **d**, Representative negative stained micrographs of the hZnT7-Fab complexes. Only the complexes with Fab#1, 3, 4, and 5 were analyzed.

**Supplementary Fig. 2.**
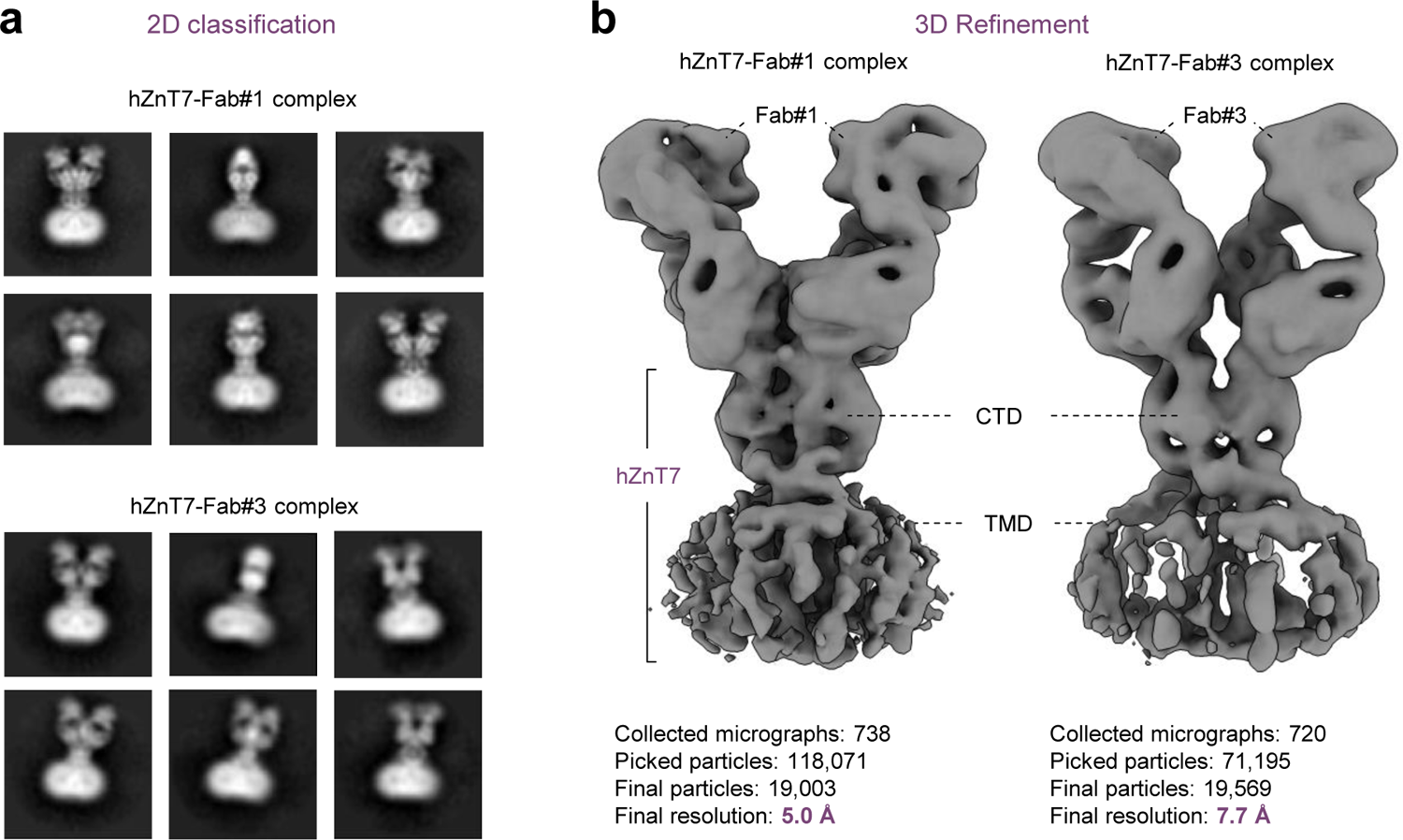
Cryo-EM density maps of hZnT7-Fab#1 and -Fab#3 complexes measured by Talos Arctica TEM. **a**, Average 2D classification of the hZnT7-Fab#1 (top) and hZnT7-Fab#3 (bottom) complexes. **b**, Cryo-EM density maps of the hZnT7-Fab#1 complex (left) at 5.0 Å-resolution and of the hZnT7-Fab#3 complex (right) at 7.7 Å-resolution, both shown at contour levels of 0.025σ in ChimeraX. TMD, transmembrane domain; CTD, cytosolic domain.

**Supplementary Fig. 3.**
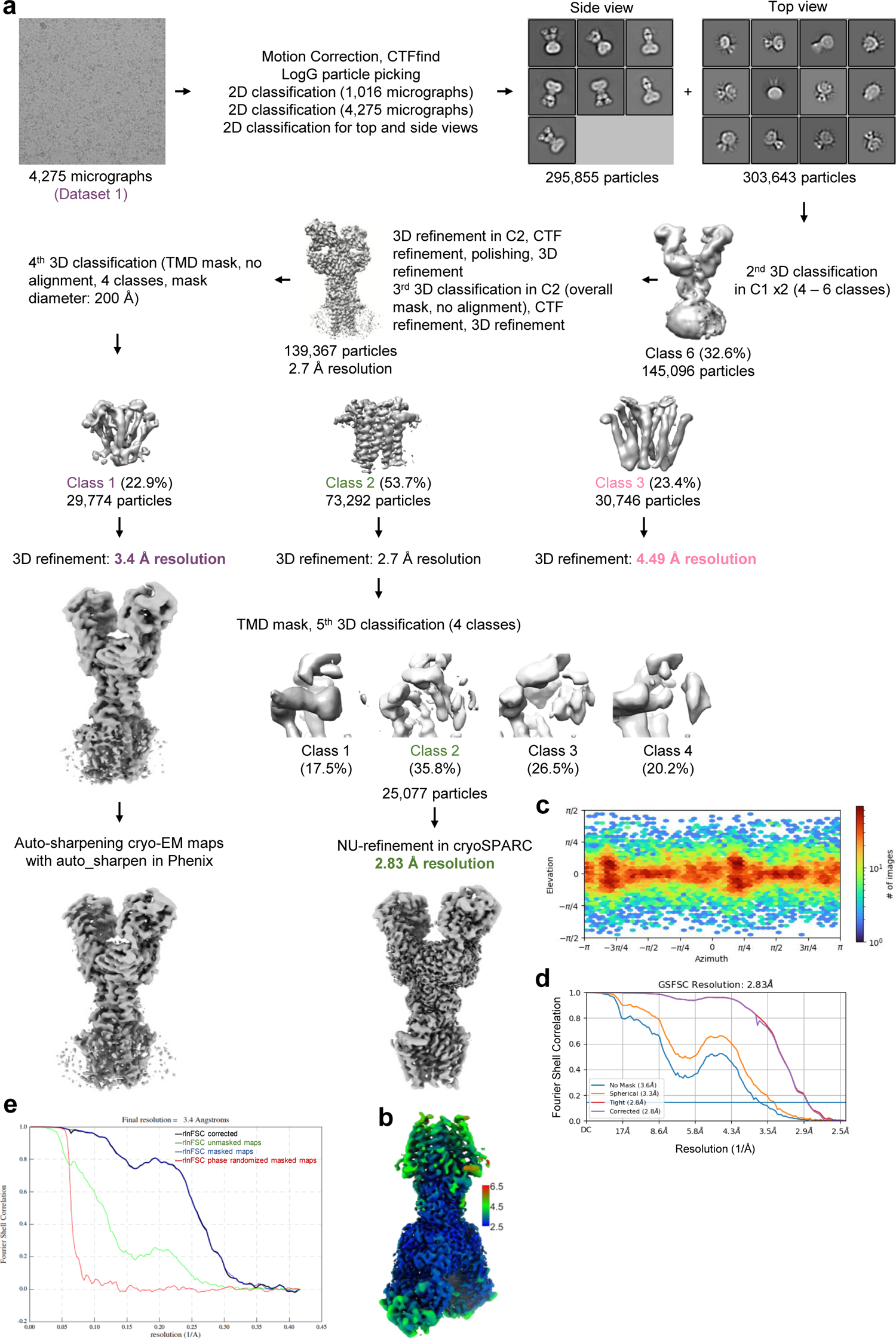
Cryo-EM analysis of the hZnT7-Fab#1 complex in the absence of zinc ions. **a**, Workflow of cryo-EM data processing and map refinement for the hZnT7-Fab#1 complex. Details are provided in the Methods section. **b**, Local-resolution map of the hZnT7-Fab#1 complex in Class 2 calculated by cryoSPARC. Bars on the right indicate local resolution in Å. **c**, Euler angle distribution of the refined particle subset used in final cryo-EM reconstruction of the hZnT7-Fab#1 complex in Class 2 by non-uniform refinement. **d**, Fourier shell correlation (FCS) plots of the hZnT7-Fab#1 complex in Class 2 calculated by cryoSPARC. **e**, Fourier shell correlation (FCS) plots of the hZnT7-Fab#1 complex in Class 1 calculated by RELION.

**Supplementary Fig. 4.**
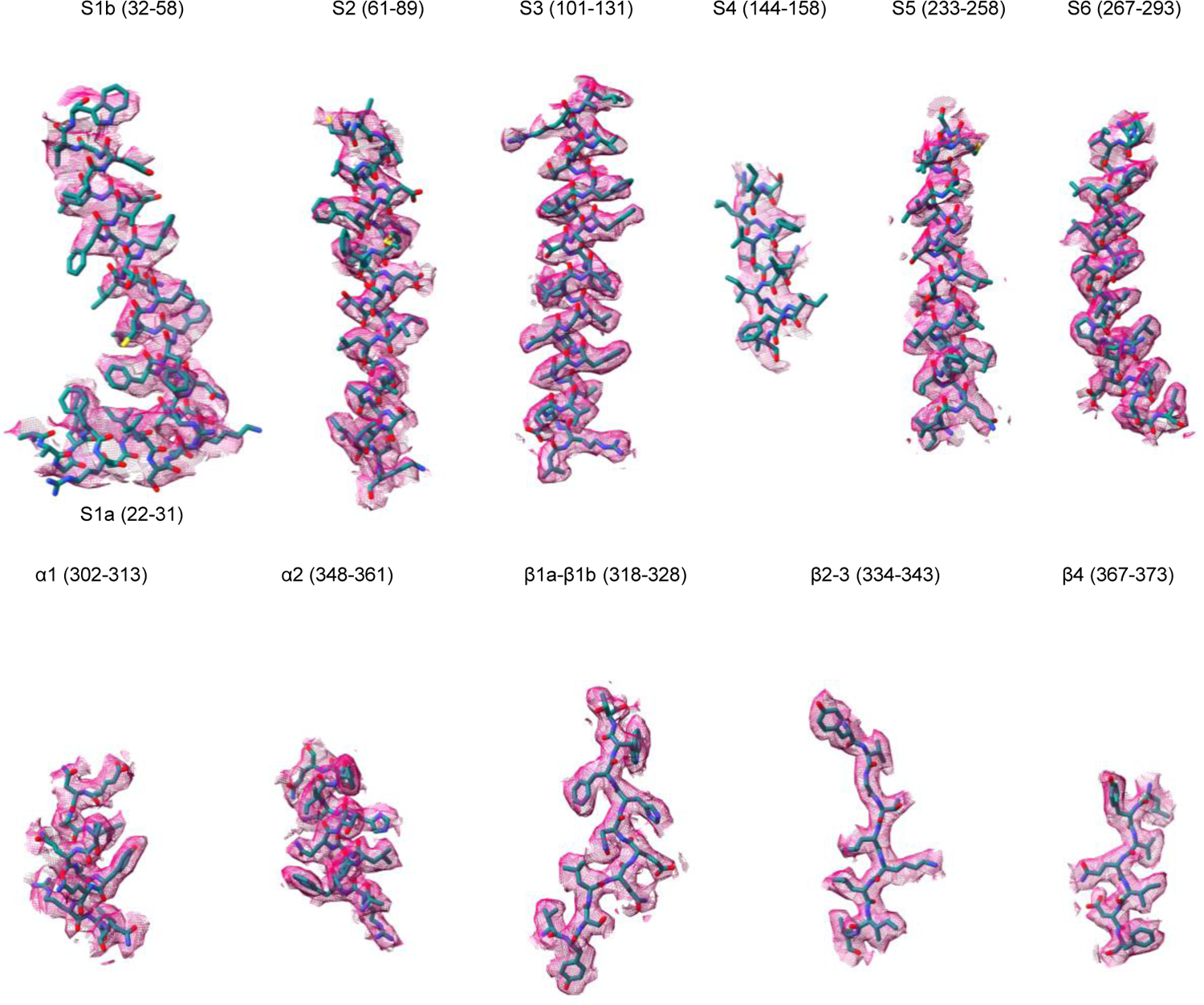
Cryo-EM density maps of primary and secondary structural elements of the OF form of Zn^2+^-unbound hZnT7. Density maps of the six transmembrane helices (S1a-S6) and cytosolic-domain elements (α1 and α2, β1a-β4) of hZnT7, with atomic models shown as sticks. All density maps are shown at a contour level of 0.5σ.

**Supplementary Fig. 5.**
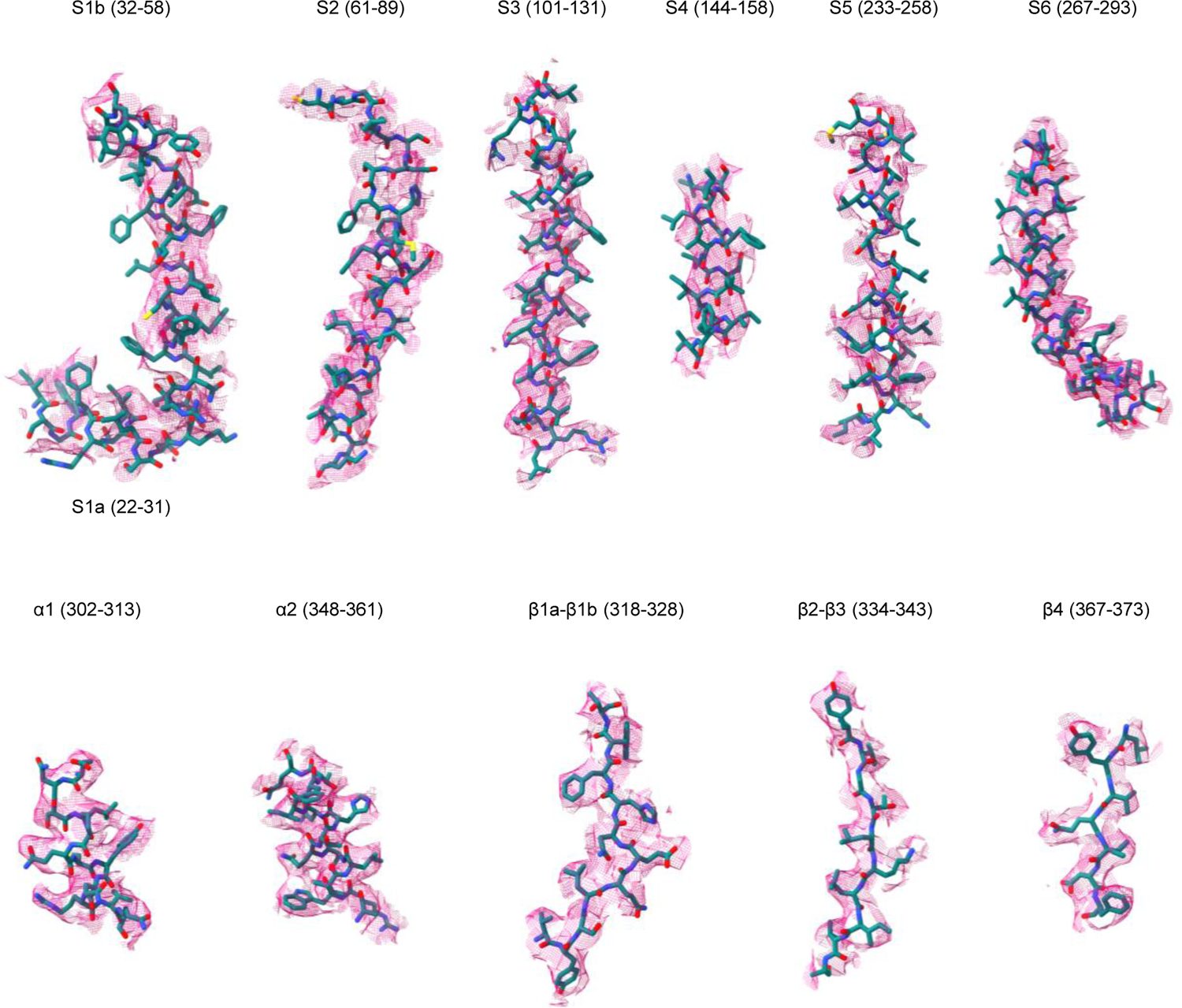
Cryo-EM density maps of primary and secondary structural elements of the IF form of Zn^2+^-unbound hZnT7. Density maps of the six transmembrane helices (S1a-S6) and cytosolic-domain elements (α1 and α2, β1a-β3) of hZnT7, with atomic models shown as sticks. All density maps are shown at a contour level of 7σ.

**Supplementary Fig. 6.**
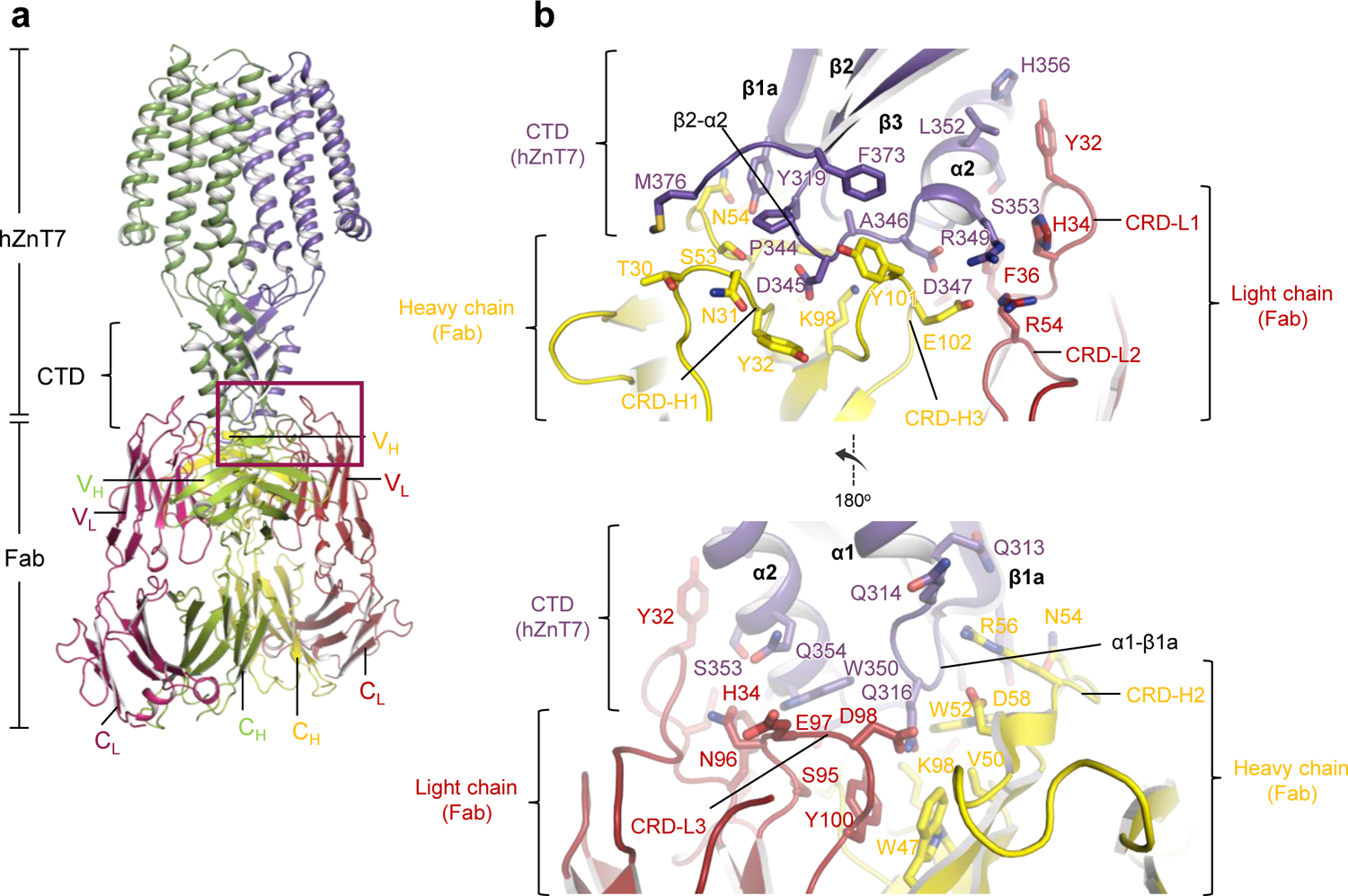
Mode of interactions between the cytosolic domain (CTD) and Fab. **a**, Cartoon representation of the Zn^2+^-unbound OF-form of hZnT7 complexed with Fab. hZnT7, Fab heavy and light chains are colored blue, yellow and beige, respectively. The interface between the CTD and Fab is marked by a red square, which is highlighted in **b**. **b**, The CTD-Fab interface. Residues involved in the CTD-Fab interactions are represented by sticks.

**Supplementary Fig. 7.**
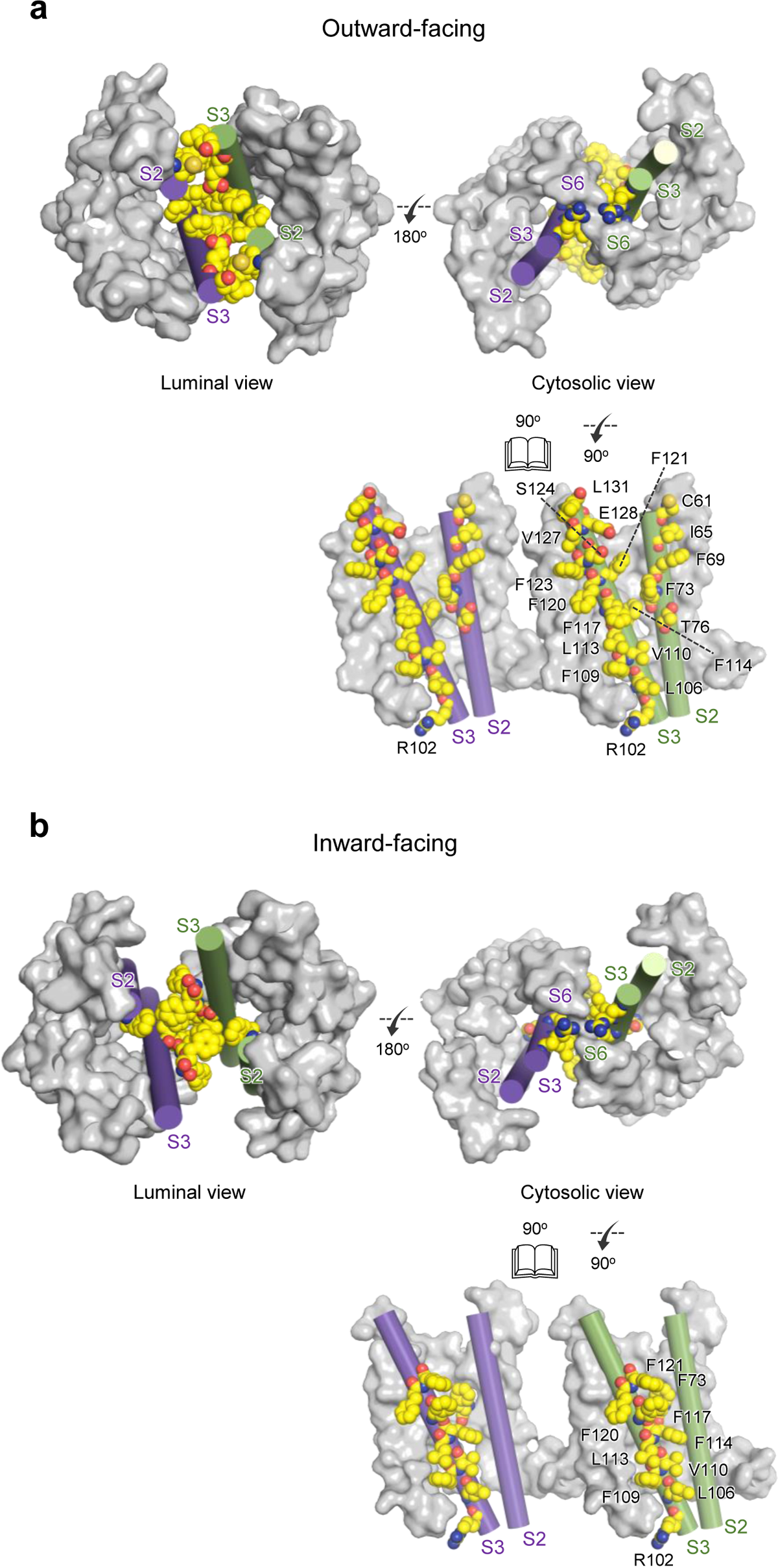
Mode of interactions between the TMDs in the OF (a) and IF (b) forms of hZnT7 homodimers. Residues involved in the interactions between S2 and S3 in the OF (a) and IF (b) forms of hZnT7 are represented by spheres. The lower figure in each panel shows a spread image of the dimer interface. S2 and S3 are shown as cylinders, while the other TM helices are shown as gray surfaces. The CTD and extra-membrane loops have been omitted for clarity.

**Supplementary Fig. 8.**
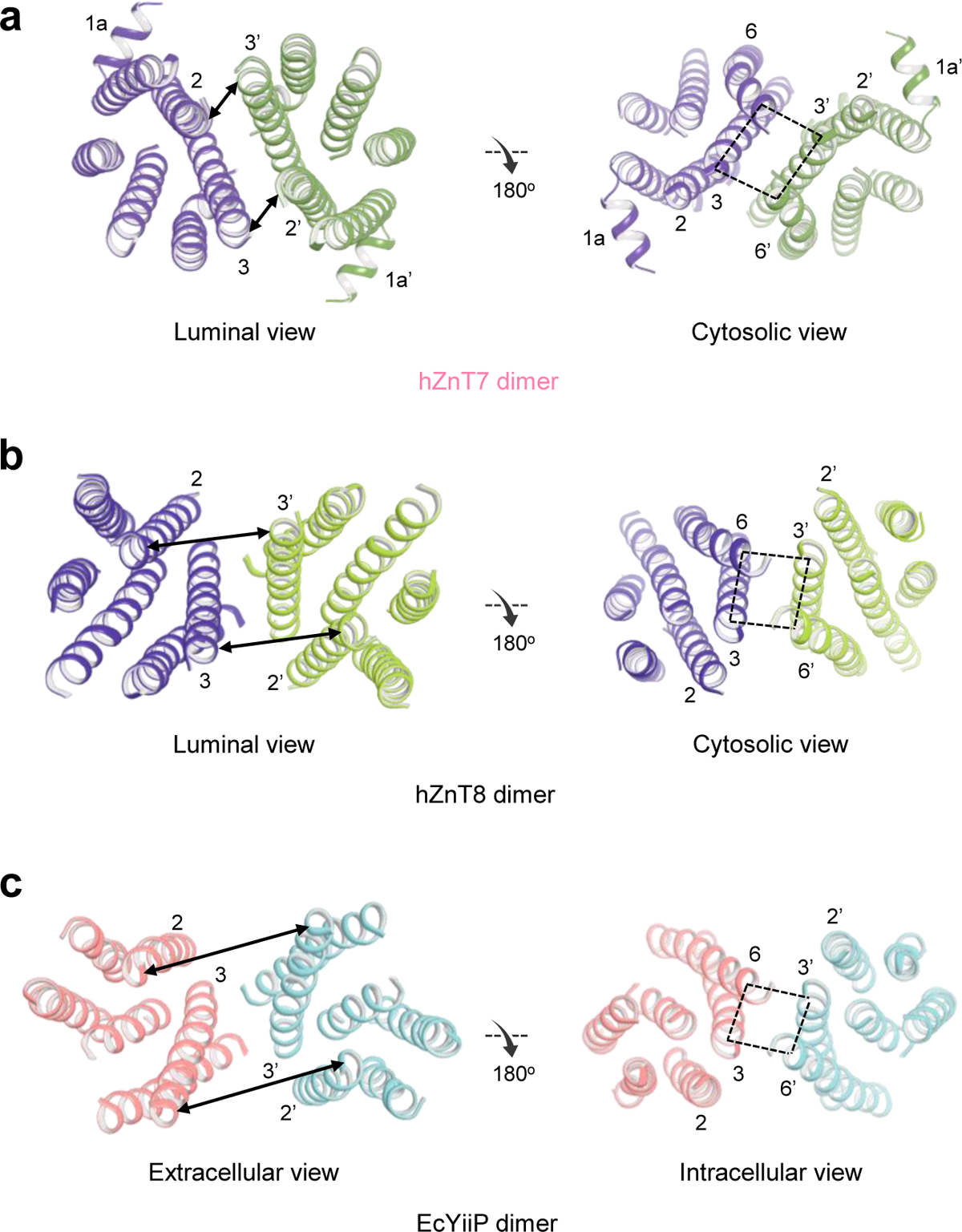
Comparison of the structures of the dimeric interfaces and transmembrane helix locations of hZnT7 (a), hZnT8 (b, PDB ID: 6XPD), and EcYiiP (c, PDB ID: 3H90). The CTD and extra-membrane loops have been omitted for clarity. Numbers indicate transmembrane helices involved in dimerization.

**Supplementary Fig. 9.**
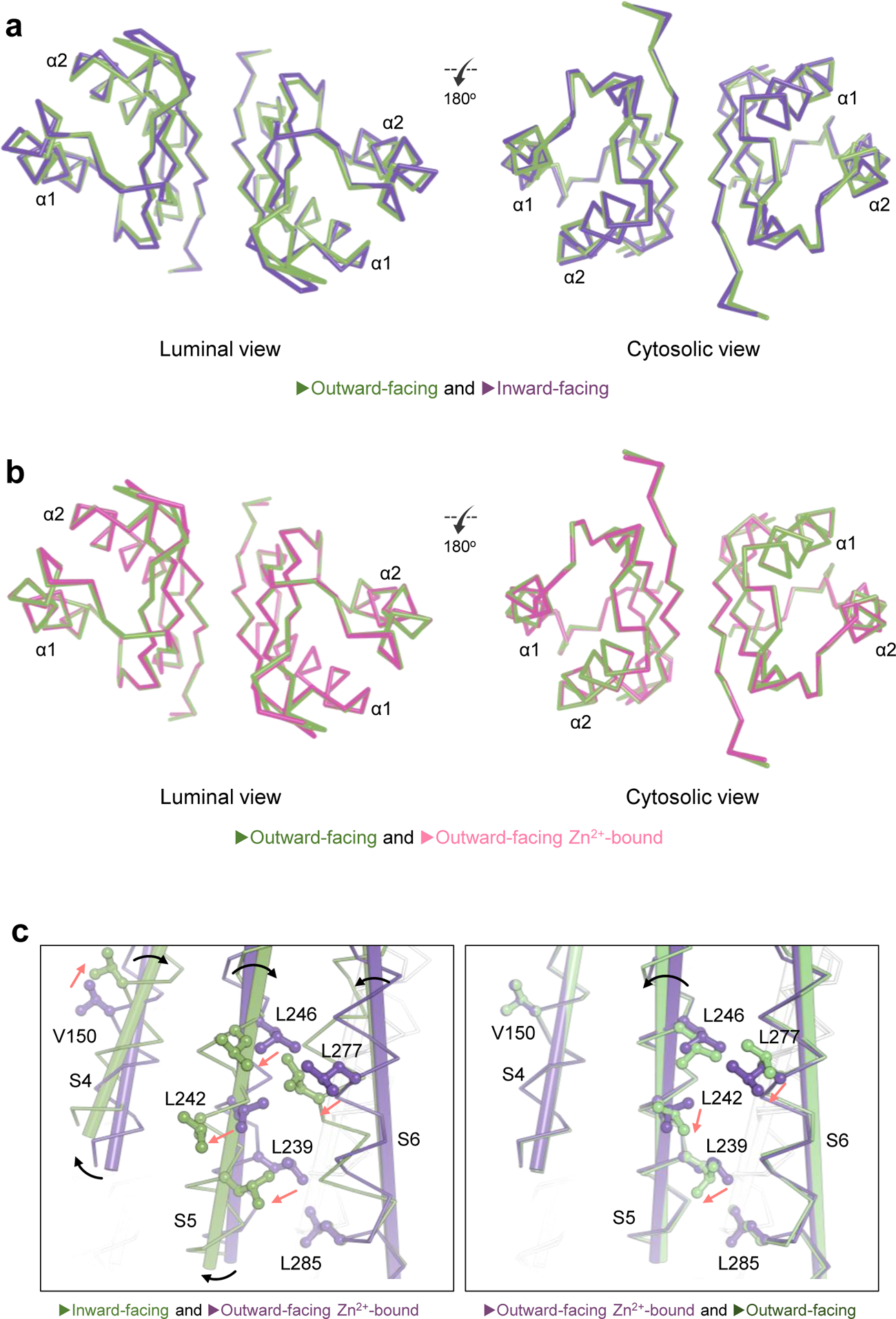
Comparison of the structures of the CTD in three different forms of hZnT7. **a**, Superimposition of CTDs of the Zn^2+^-unbound OF (purple) and IF (green) forms of hZnT7. **b**, Superimposition of CTDs of the OF forms of Zn^2+^-unbound (green) and -bound (pink) hZnT7. **c**. Comparison of the structures of the TMD in three different forms of hZnT7 by superimposition of their CTDs. Leucine and valine residues are shown as sticks. Black arrows indicate the movement of the TM helices, and red arrows indicate the movement of amino acid residues.

**Supplementary Fig. 10.**
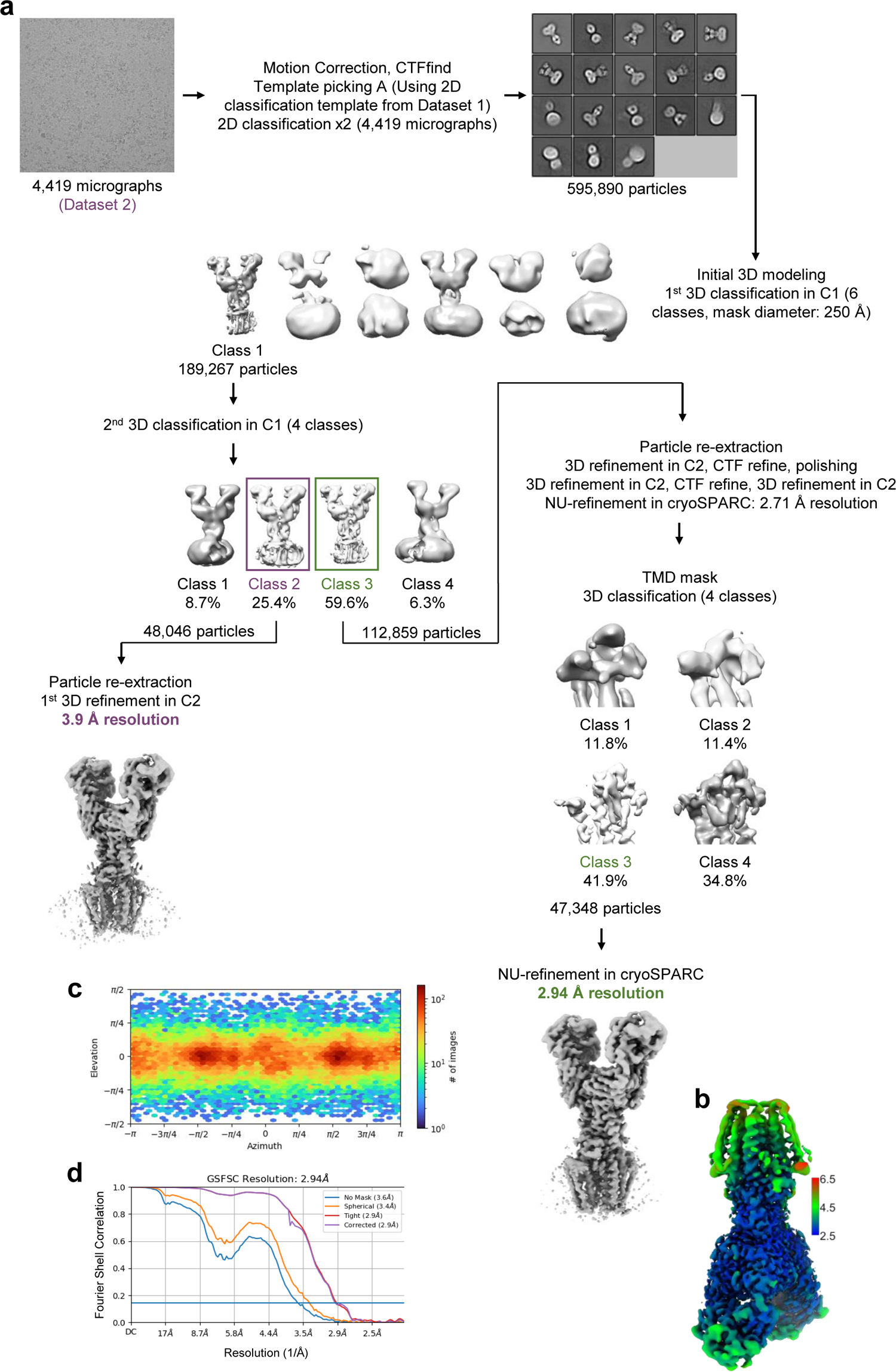
Cryo-EM analysis of Zn^2+^-bound hZnT7 complexed with Fab#1. **a**, Workflow of cryo-EM data processing and map refinement for the hZnT7 (Zn^2+^-bound)-Fab#1 complex. Details are provided in the Methods section. **b**, Local-resolution map of the hZnT7-Fab#1 complex in Class 3 calculated by cryoSPARC. Bars on the right indicate local resolution in Å. **c**, Euler angle distribution of the refined particle subset used in final cryo-EM reconstruction of the hZnT7-Fab#1 complex in Class 3 by non-uniform refinement. **d**, Fourier shell correlation (FCS) plots of the hZnT7-Fab#1 complex in Class 3 calculated by cryoSPARC.

**Supplementary Fig. 11.**
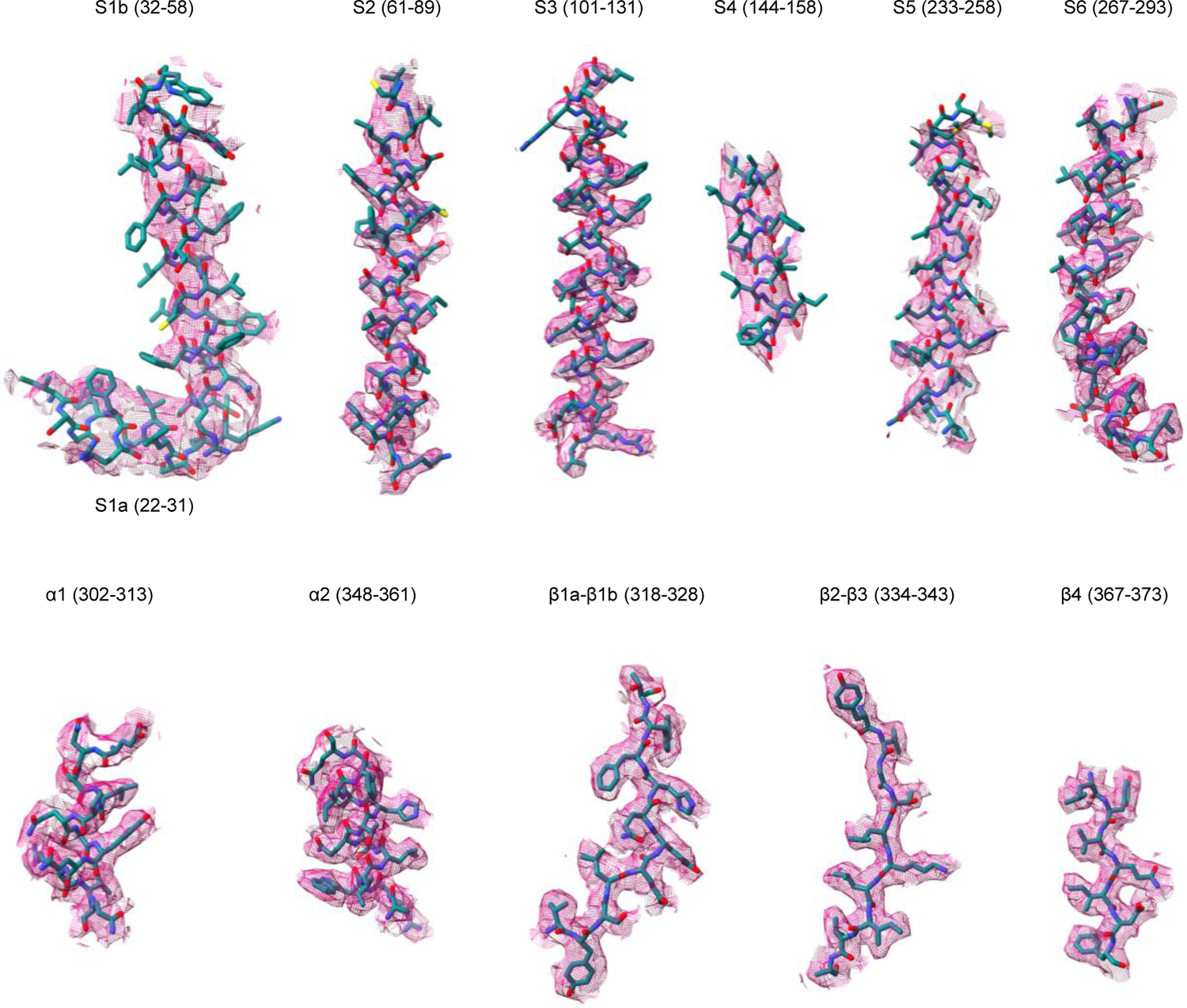
Cryo-EM density maps of primary and secondary structural elements of the OF form of Zn^2+^-bound hZnT7. Density maps of the six transmembrane helices (S1a-S6) and cytosolic-domain elements (α1 and α2, β1a-β3) of hZnT7 with atomic models shown as sticks. All density maps are shown at a contour level of 0.5σ.

**Supplementary Fig. 12.**
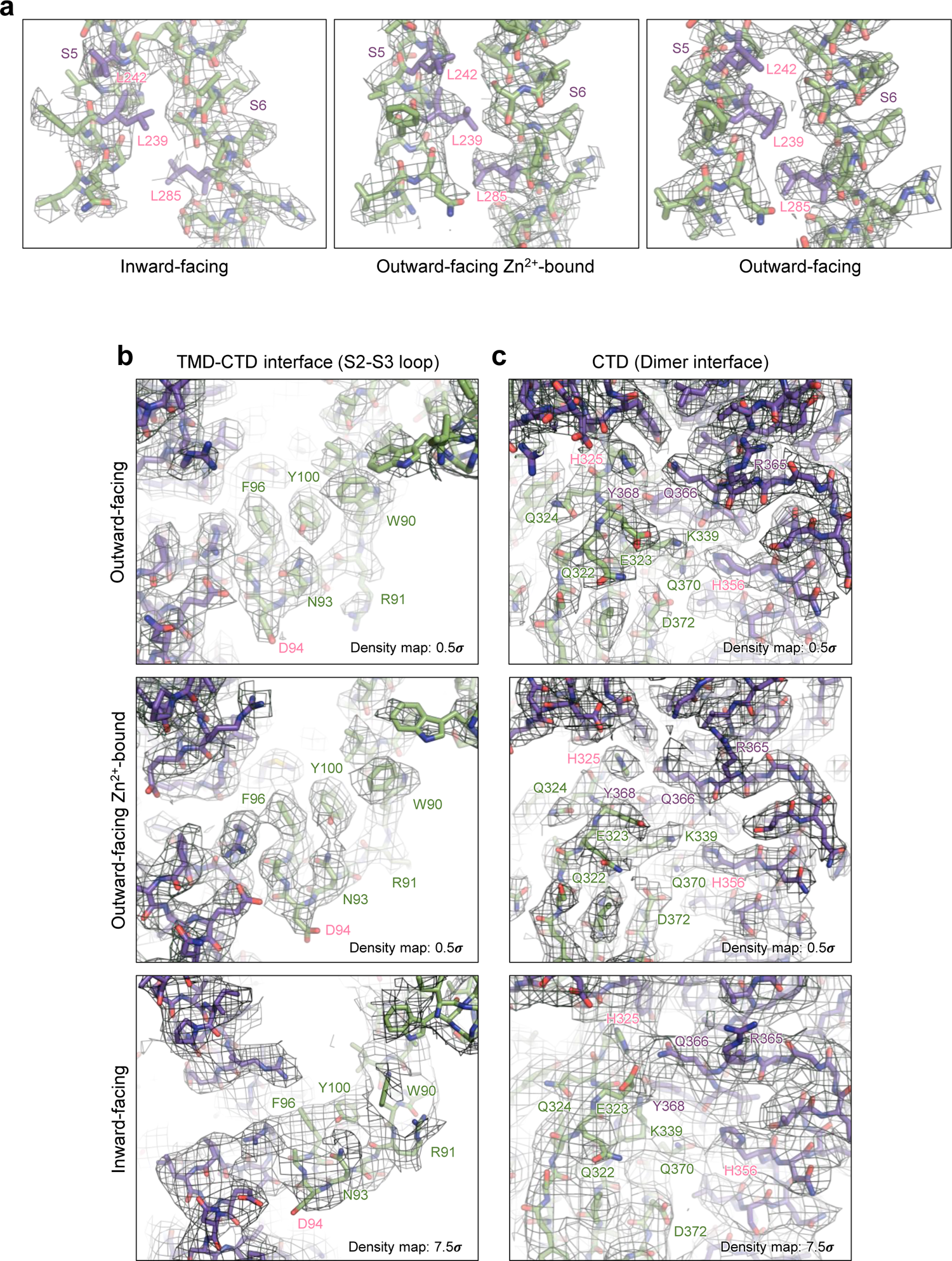
Cryo-EM maps of the leucine-pair gate in hZnT7. (a), the S2-S3 loop at the TMD-CTS interface (b), and the dimer interface in the CTD (c) of hZnT7. a, Cryo-EM density maps of the Leu^239^ (S5)- and Leu^285^ (S6)-neighboring regions in the indicated states are displayed as green mesh. Leu^239^, Leu^242^, and Leu^285^ are colored violet. Cryo-EM density maps are shown at contour levels of 7.0σ (inward-facing) and 0.4σ (outward-facing and outward-facing Zn^2+^-bound). b, Cryo-EM density maps of the S2-S3 loop at the TMD-CTD interface in the indicated states are displayed as green mesh. c, Cryo-EM density maps of the dimer interface in the CTD in the indicated states are represented by green (protomer 1) and violet (protomer 2) meshes. Cryo-EM density maps are shown at contour levels of 0.5σ (outward-facing and outward-facing Zn^2+^-bound) and 7.5σ (inward-facing). Protein models are shown as sticks.

**Supplementary Fig. 13.**
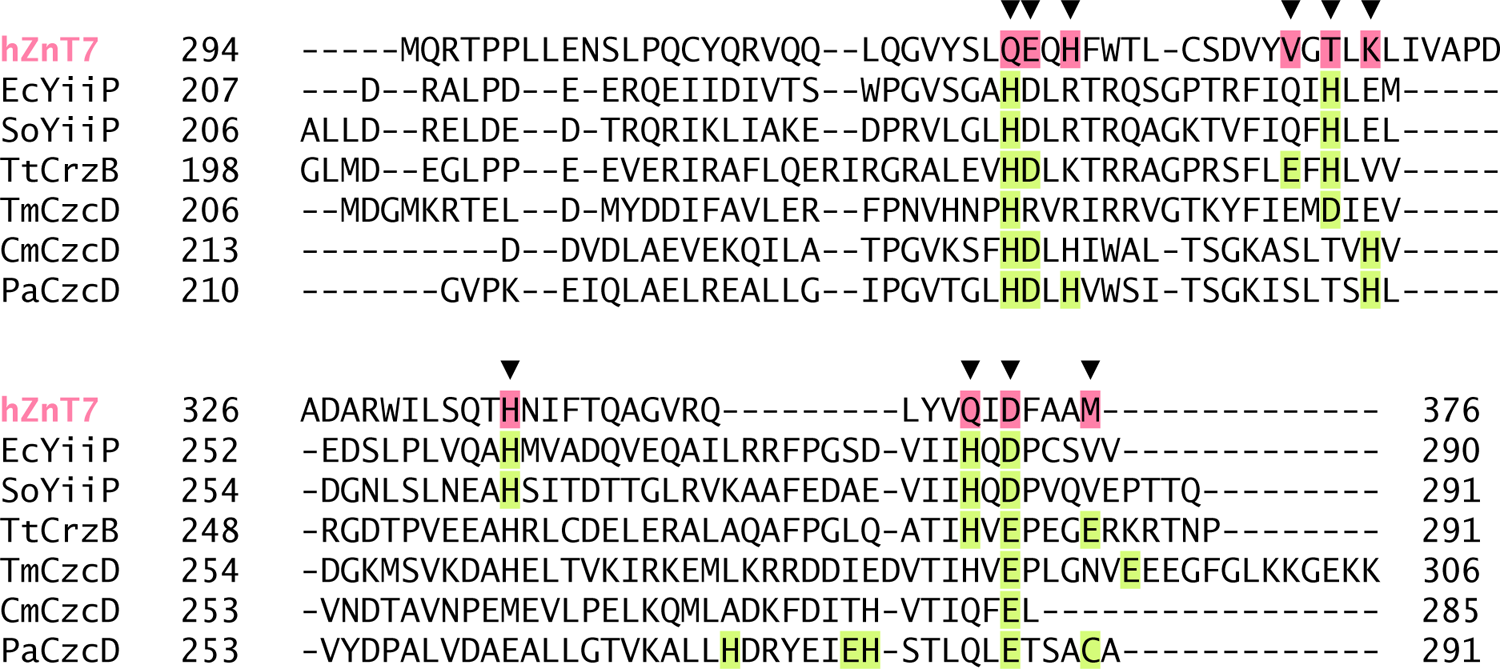
Comparison of Zn^2+^-binding residues in hZnT7 and other zinc transporters. Amino acid sequence alignment of the C-terminal regions of hZnT7 and bacterial Zn^2+^ transporters. The C-terminal segment sequences were downloaded from the UniProt site for human ZnT7 (hZnT7) (UniProtKB: Q8NEW0), *Escherichia coli* YiiP (EcYiiP) (UniProtKB: P69380), *Shewanella oneidensis* YiiP (SoYiiP) (UniProtKB: Q8E919), *Thermus thermophilus* CzrB (TtCzrB) (UniProtKB: Q8VLX7), *Thermotoga maritima* CzcD (TmCzcD) (UniProtKB: Q9WZX9), *Cupriavidus metallidurans* CzcD (CmCzcD) (UniProtKB: P13512), and *Pseudomonas aeruginosa* CzcD (PaCzcD) (UniProtKB: Q9I6A3). Residues involved in Zn^2+^ binding are indicated by black arrowheads and marked by green boxes. The corresponding residues in hZnT7 are colored pink.

**Supplementary Fig. 14.**
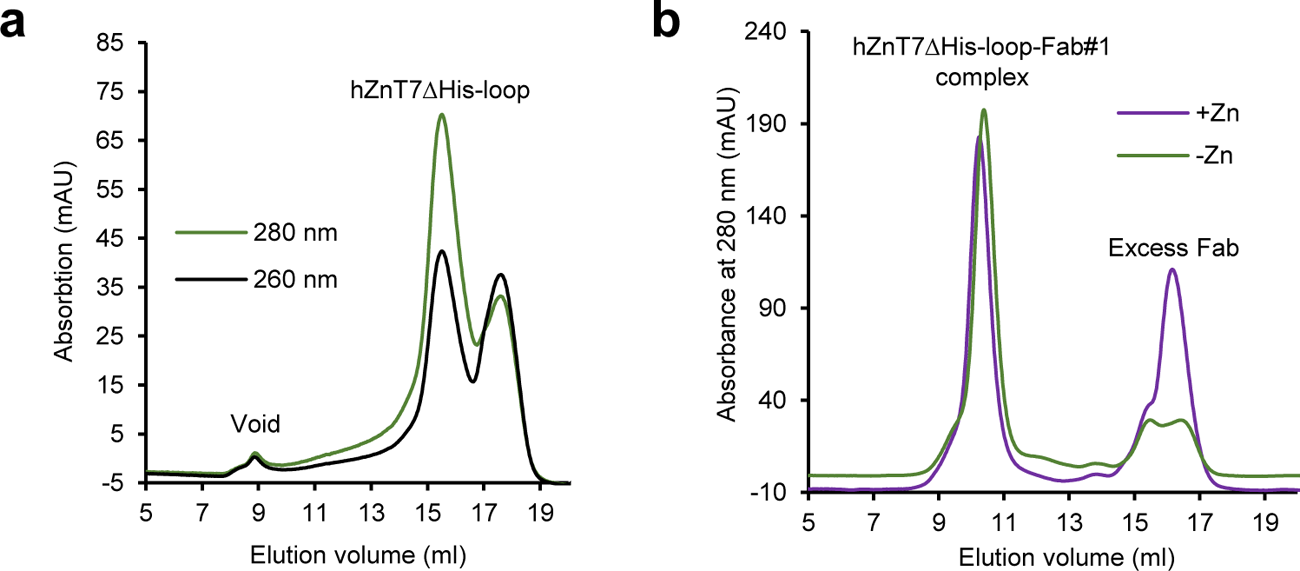
Purification and preparation of the hZnT7ΔHis-loop-Fab#1 complex for cryo-EM analysis. Representative chromatograms from **a**, Superose 6 Increase gel filtration of hZnT7ΔHis-loop protein and **b**, Superdex 200 gel filtration of the hZnT7ΔHis-loop-Fab#1 complex without (green) and with (purple) 10 μM ZnSO_4_.

**Supplementary Fig. 15.**
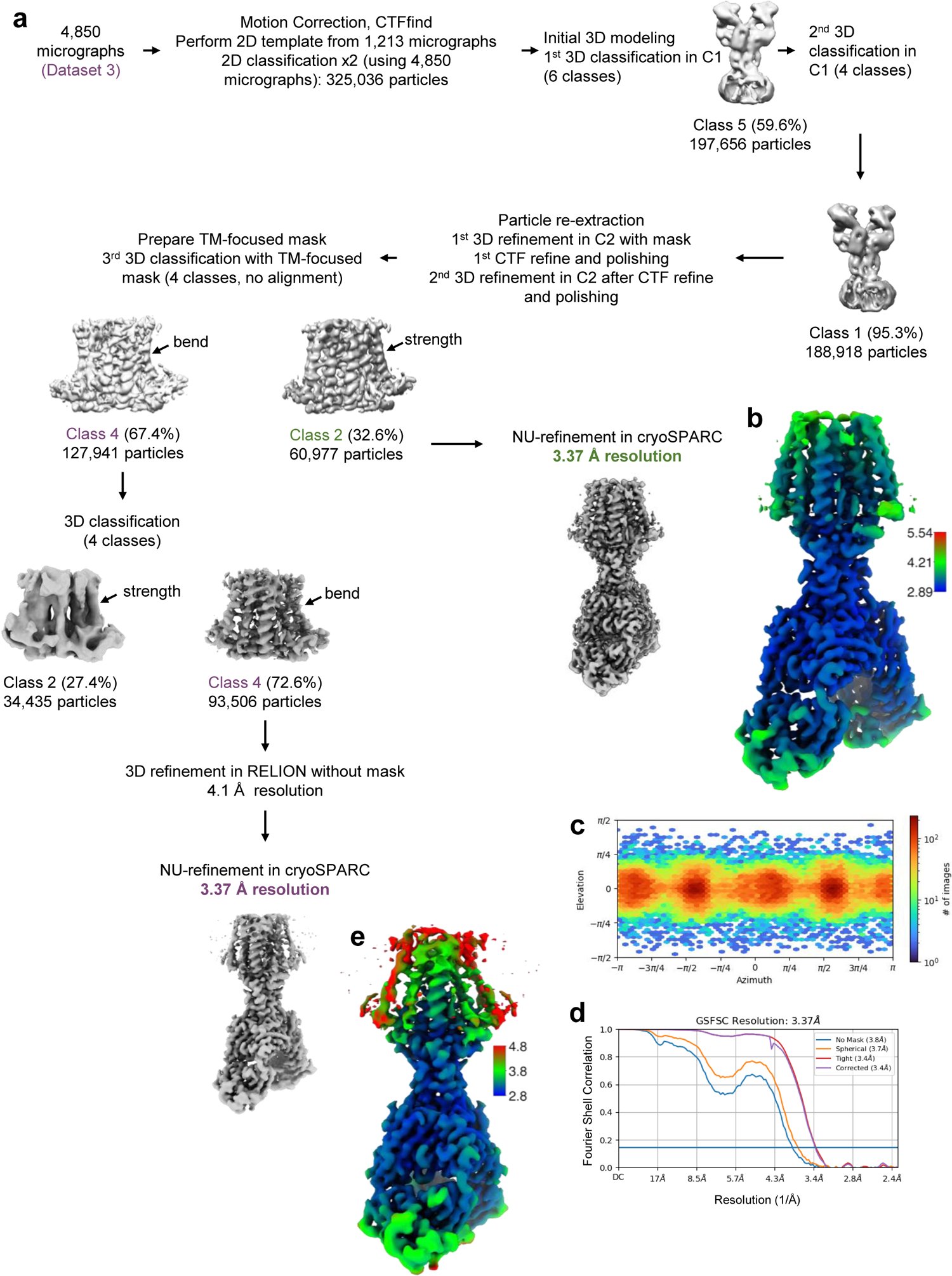
Cryo-EM analysis of Zn^2+^-unbound hZnT7ΔHis-loop in complex with Fab#1. **a**, Workflow of cryo-EM data processing and map refinement of the hZnT7ΔHis-loop (Zn^2+^-unbound)-Fab#1 complex. **b** and **e**, Local-resolution map of the hZnT7ΔHis-loop-Fab#1 complex in Class 2 (b) and Class 4 (e) calculated by cryoSPARC. Bars on the right indicate local resolution in Å. **c**, Euler angle distribution of the refined particle subset used in final cryo-EM reconstruction of the hZnT7ΔHis-loop (Zn^2+^-unbound)-Fab#1 complex by non-uniform refinement. **d**, Fourier shell correlation (FCS) plots of the hZnT7ΔHis-loop-Fab#1 complex in Class 2.

**Supplementary Fig. 16.**
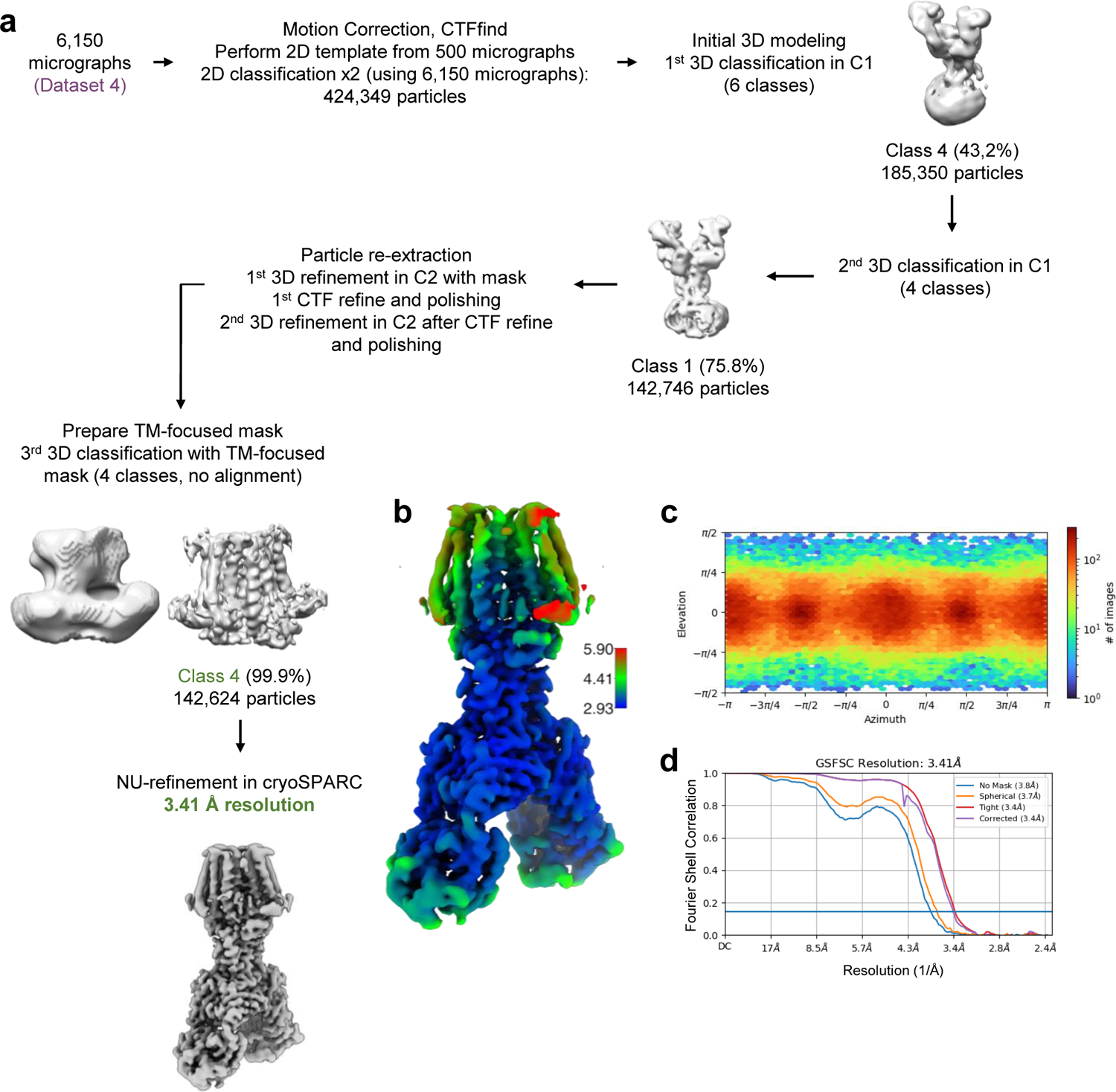
Cryo-EM analysis of Zn^2+^-bound hZnT7ΔHis-loop in complex with Fab#1. **a**, Workflow of cryo-EM data processing and map refinement of the hZnT7ΔHis-loop (Zn^2+^-bound)-Fab#1 complex. **b**, Local-resolution map of the hZnT7ΔHis-loop-Fab#1 complex in Class 4 calculated by cryoSPARC. Bars on the right indicate local resolution in Å. **c**, Euler angle distribution of refined particle subset used in final cryo-EM reconstruction of the hZnT7ΔHis-loop(Zn^2+^-bound)-Fab#1 complex by non-uniform refinement. **d**, Fourier shell correlation (FCS) plot of the hZnT7ΔHis-loop (Zn^2+^-bound)-Fab#1 complex in Class 4.

**Supplementary Fig. 17.**
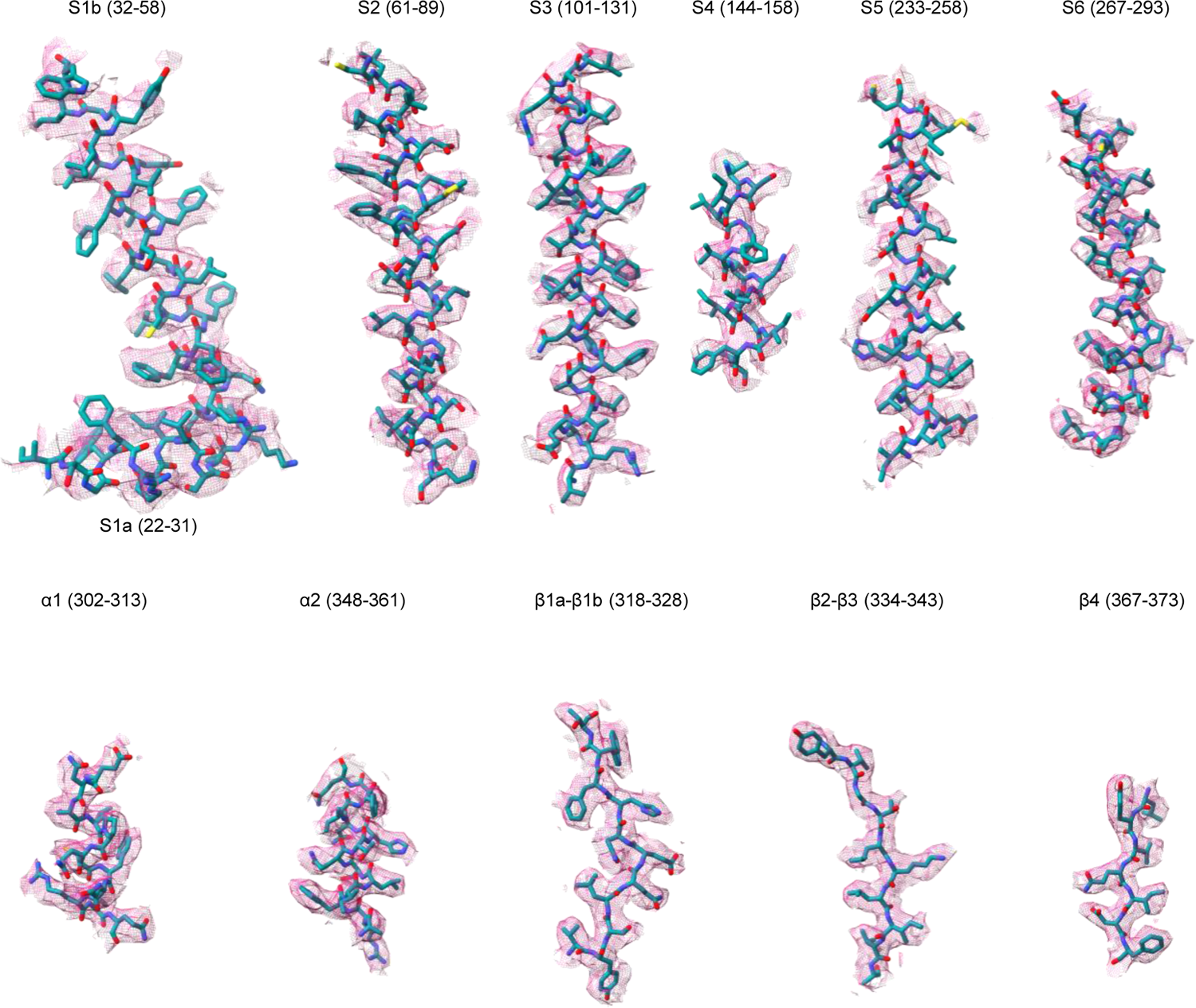
Cryo-EM density maps of primary and secondary structural elements of the OF form of Zn^2+^-unbound hZnT7ΔHis-loop. Density maps of the six transmembrane helices (S1a-S6) and cytosolic-domain elements (α1 and α2, β1a-β4) of Zn^2+^-unbound hZnT7ΔHis-loop with their atomic models indicated with sticks. All density maps are shown at contour levels of 0.5σ.

**Supplementary Fig. 18.**
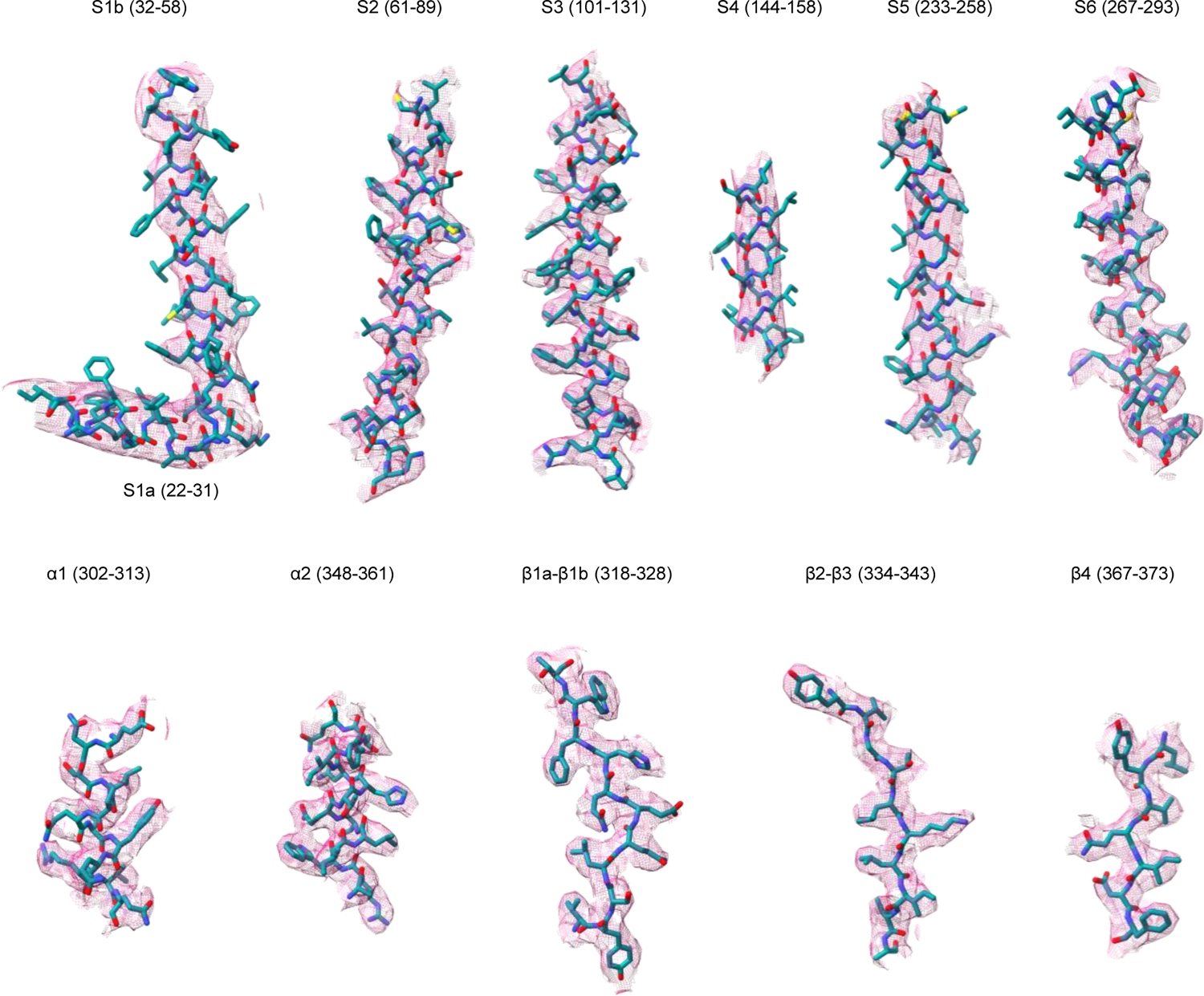
Cryo-EM density maps of primary and secondary structural elements of the OF form of Zn^2+^-bound hZnT7ΔHis-loop. Density maps of the six transmembrane helices (S1a-S6) and cytosolic-domain elements (α1 and α2, β1a-β4) of Zn^2+^-bound hZnT7ΔHis-loop with their atomic models indicated with sticks. All density maps are shown at contour levels of 0.5σ.

**Supplementary Fig. 19.**
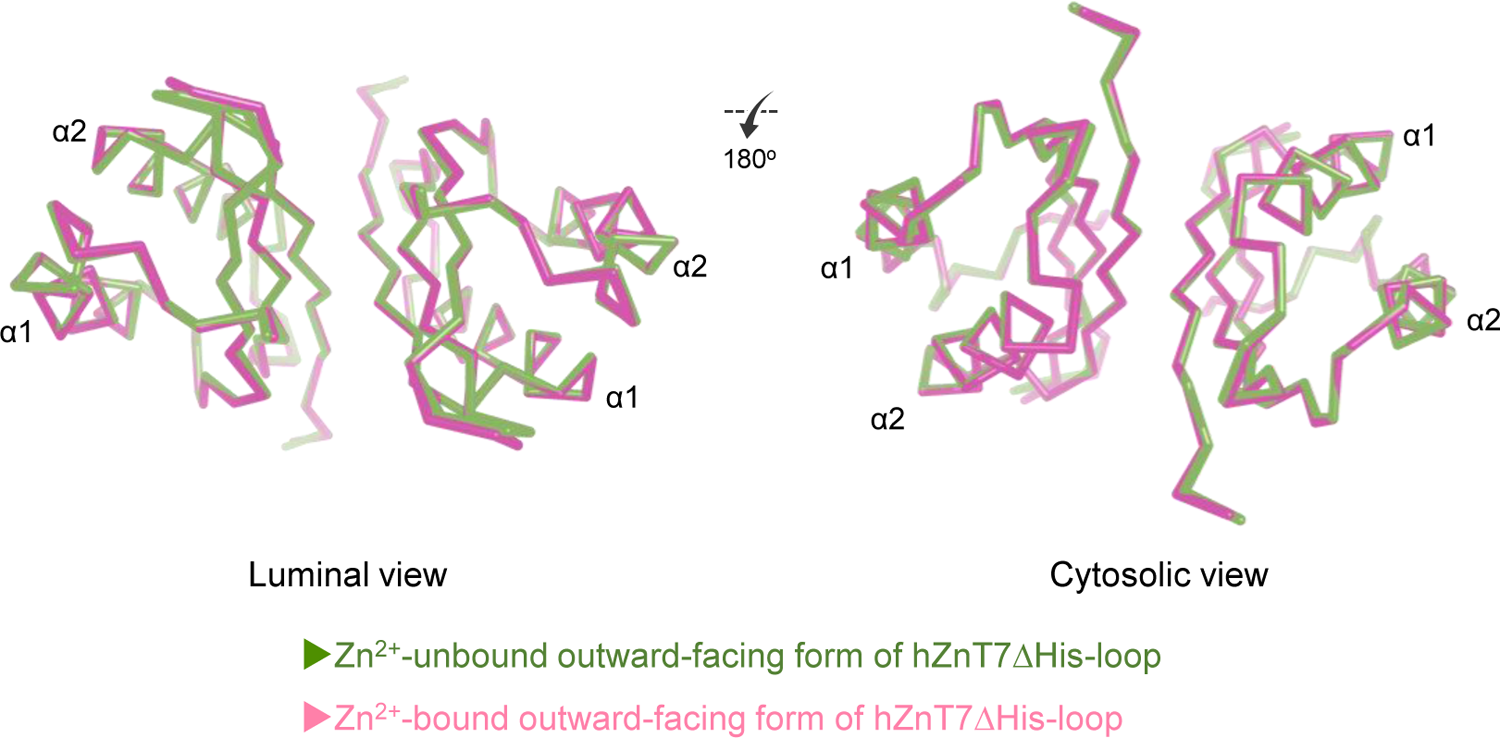
Structural comparison of the CTDs of Zn^2+^-unbound and -bound forms of hZnT7ΔHis-loop.

**Supplementary Fig. 20.**
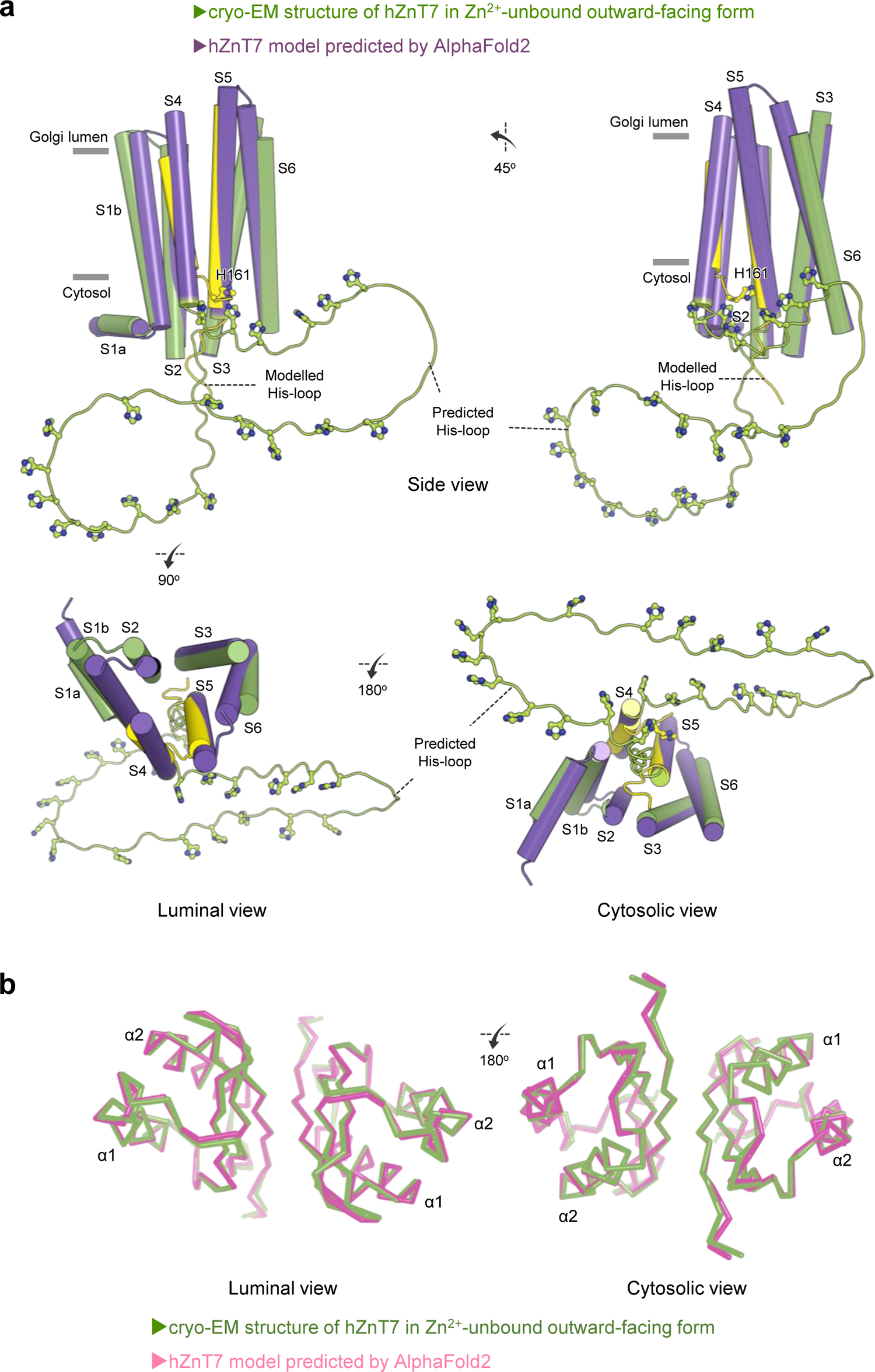
Predicted structural model of hZnT7. a, Entire structure of hZnT7 predicted by AlphaFold2. S4 and S5 are highlighted in yellow. Histidine residues on the putative His-loop model are indicated by green sticks. All transmembrane helices are represented by cylinders. b, Superimposition of the CTD of the cryo-EM structures of Zn^2+^-unbound hZnT7 (green) and the hZnT7 (pink) model predicted by AlphaFold2.

## Supplementary

Amino acid sequence of Fab#1 (YN7114-08)

### Light chain

>YN7114-08 Fab light chain_aa seq DIVLTQSPASLAVSLRRRATISCRASESVDGYGHSFMHWYQQKSGQPPKLLIYRASNLESGVPARFSGSGSRTDFTLTIDPVEADDAATYYCQQSNEDPYTFGSGTKLEIKRADAAPTVSIFPPSSEQLTSGGASVVCFLNNFYPKDINVKWKIDGSERQNGVLNSWTDQDSKDSTYSMSSTLTLTKDEYERHNSYTCEATHKTSTSPIVKSFNRNEC

CDR-L1: RASESVDGYGHSFMH

CDR-L2: ASNLESG

CDR-L3: QQSNEDPYT

V_L_ region: DIVLTQ………TKLEIK

C_L_ region: RADAAPT………SFNRNEC

### Heavy chain

>YN7114-08 Fab heavy chain_aa seq

EVQLQESGPGLVAPSQSLSITCTVSGFSLTNYAVHWVRQSPGKGLEWLGVIWSNGRTDYNAAFISRLSISKDNSKSQVFFKMNSLQADDTAIYYCARKLAYEGAMDYWGQGTSVTVSSAKTTPPSVYPLAPGSAAQTNSMVTLGCLVKGYFPEPVTVTWNSGSLSSGVHTFPAVLQSDLYTLSSSVTVPSSTWPSETVTCNVAHPASSTKVDKKIVPRDCGCKPCICTVPEVSS

CDR-H1: NYAVH

CDR-H2: VIWSNGRTDYNAAFIS

CDR-H3: KLAYEGAMDY

V_H_ region: EVQ………TSVTVSS

C_H_ region: AKTTPPS………CICTVPEVSS

## References

1. Andreini, C., Banci, L., Bertini, I. & Rosato, A. Counting the zinc-proteins encoded in the human genome. J. Proteome. Res. 5:196–201 (2006).

2. Eide, D. J. Zinc transporters and the cellular trafficking of zinc. Biochim. Biophys. Acta. 1763:711–722 (2006).

3. Kambe, T. Molecular architecture and function of ZnT transporters. Curr. Top. Membr. 69:199–220 (2012).

4. Kambe, T., Taylor, K. M. & Fu, D. Zinc transporters and their functional integration in mammalian cells. J. Biol. Chem. 296:100320 (2021).

5. Gao, H.L., Feng, W.Y., Li, X.L., Xu, H., Huang, L.P. & Wang, Z.Y. Golgi apparatus localization of ZNT7 in the mouse cerebellum. Histol. Histopathol. 24:567–572 (2009).

6. Chi, Z.H., Feng, W.Y., Gao, H.L., Zheng, W., Huang, L. & Wang, Z.Y. ZNT7 and Zn^2+^ are present in different cell populations in the mouse testis. Histol. Histopathol. 24:25–30 (2009).

7. Kirschke, C. P. & Huang, L. P. ZnT7, a novel mammalian zinc transporter, accumulates zinc in the Golgi apparatus. J. Biol. Chem. 278:4096–4102 (2003).

8. Gao, H.L., Xu, H., Wang, X., Dahlstrom, A., Huang, L.P. & Wang, Z.Y. Expression of zinc transporter ZnT7 in mouse superior cervical ganglion. Auton Neurosci. 140:59–65 (2008).

9. Kowada, T., Watanabe, T., Amagai, Y., Liu, R., Yamada, M., Takahashi, H., Inaba, K. & Mizukami, S. Quantitative imaging of labile Zn^2+^ in the Golgi apparatus using a localizable small-molecule fluorescent probe. Cell Chem. Biol. 27:1521–1531 (2020).

10. Amagai, Y. et al. Zinc homeostasis governed by Golgi-resident ZnT family members regulates ERp44-mediated proteostasis in the early secretory pathway. In revision (2022).

11. Huang, L. P., Yu, Y. Y., Kirschke, C. P., Gertz, E. R. & Lloyd, K. K. C. ZnT7 (Slc30a7)-deficient mice display reduced body zinc status and body fat accumulation. J. Biol. Chem. 82:37053–37063 (2007).

12. Tepaamorndech, S., Huang, L. P. & Kirschke, C. P. A null-mutation in the Znt7 gene accelerates prostate tumor formation in a transgenic adenocarcinoma mouse prostate model. Cancer Lett. 308:33–42 (2011).

13. Tepaamorndech, S., Kirschke, C. P., Pedersen, T. L., Keyes, W. R., Newman, J. W. & Huang, L. P. Zinc transporter 7 deficiency affects lipid synthesis in adipocytes by inhibiting insulin-dependent Akt activation and glucose uptake. FEBS J. 283:378–94 (2016).

14. Lu, M. & Fu, D. Structure of the zinc transporter YiiP. Science 317:1746–1748 (2007).

15. Lu, M., Chai, J. & Fu, D. Structural basis for autoregulation of the zinc transporter YiiP. Nat. Struct. Mol. Biol. 16:1063–7 (2009).

16. Coudray, N., Valvo, S., Hu, M., Lasala, R., Kim, C., Vink, M., Zhou, M., Provasi, D., Filizola, M., J. Tao, J., Fang, J., Penczek, A. P., Ubarretxena-Belandia, I. & Stokes, D. L. Inward-facing conformation of the zinc transporter YiiP revealed by cryoelectron microscopy. Proc. Natl. Acad. Sci. 110:2140–2145 (2013).

17. Lopez-Redondo, M. L., Coudray, N., Zhang, Z., Alexopoulos, J. & Stokes D. L. Structural basis for the alternating access mechanism of the cation diffusion facilitator YiiP. Proc. Natl. Acad. Sci. 115:3042–3047 (2018).

18. Lopez-Redondo, M., Fan, S. J., Koide, A., Koide, S., Beckstein, O. & Stokes, L. D. Zinc binding alters the conformational dynamics and drives the transport cycle of the cation diffusion facilitator YiiP. J. Gen Physiol. 153:e202112873 (2021).

19. Xue, J., Xie, T., Zeng, W. Z., Jiang, Y. X. & Bai, X. C. Cryo-EM structures of human ZnT8 in both outward- and inward-facing conformations. Elife 9:e58823 (2020).

20. Gupta, S., Chai, J., Cheng, J., D’Mello, R., Chance, M. R. & Fu, D. Visualizing the kinetic power stroke that drives proton-coupled zinc(II) transport. Nature 512:101–4 (2014).

21. Jaenecke, F. et al. Generation of conformation-specific antibody fragments for crystallization of the multidrug resistance transporter MdfA. Methods Mol. Biol. 1700:97–109 (2018).

22. Scheres, S. H. RELION: implementation of a Bayesian approach to cryo-EM structure determination. J. Struct. Biol. 180:519–530 (2012).

23. Mastronarde, D. N. Automated electron microscope tomography using robust prediction of specimen movements. J. Struct. Biol. 152:36–51 (2005).

24. Rohou, A. & Grigorieff, N. CTFFIND4: Fast and accurate defocus estimation from electron micrographs. J. Struct. Biol. 192:216–221 (2015).

25. Punjani, A., Rubinstein, J. L., Fleet, D. J. & Brubaker, M. A. cryoSPARC: algorithms for rapid unsupervised cryo-EM structure determination. Nat. Methods 14:290–296 (2017).

26. Terwilliger, C. T., Sobolev, V. O., Afonine, V. P. & Adams, D. P., Automated map sharpening by maximization of detail and connectivity. Acta Crystallogr Struct Biol. 74:545–559 (2018).

27. Burnley, T., Palmer, C. M. & Winn, M. Recent developments in the CCP-EM software suite. Acta Crystallogr D Struct Biol. 73:469–477 (2017).

28. Emsley, P. & Cowtan, K. Coot: model-building tools for molecular graphics. Acta Crystallogr. 60:2126–2132 (2004).

29. Emsley, P., Lohkamp, B., Scott, W. G. & Cowtan, K. Features and development of Coot. Acta Crystallogr. 66:486–501 (2010).

30. Adams, P. D. et al. PHENIX: a comprehensive Python-based system for macromolecular structure solution. Acta Crystallogr. 66:213–221 (2010).

31. Davis, I. W. et al. MolProbity: all-atom contacts and structure validation for proteins and nucleic acids. Nucleic Acids Res. 35:W375–W383 (2007).

32. DeLano, W. L. The PyMOL molecular graphics system. http://www.pymol.org (2002).

33. Pettersen, E. F. et al. UCSF Chimera-a visualization system for exploratory research and analysis. J. Comput. Chem. 25:1605–1612 (2004).

34. Goddard, T. D. et al. UCSF ChimeraX: Meeting modern challenges in visualization and analysis. Protein Sci. 27:14–25 (2018).

35. Smart, O. S., Neduvelil, J. G., Wang, X., Wallace, B. A. & Sansom, M. S. HOLE: a program for the analysis of the pore dimensions of ion channel structural models. J. Mol. Graph. 14:354–360 (1996).

36. Humphrey, W., Dalke, A. & Schulten, K. VMD: Visual molecular dynamics. J. Mol. Graph. 14:33–38 (1996).

37. Katoh, K., Rozewicki, J. & Yamada, K. D. MAFFT online service: multiple sequence alignment, interactive sequence choice and visualization. Brief Bioinformatic 20:1160–1166 (2019).

38. Jurrus, E., Engel, D., Star, K., Monson, K., Brandi, J., Felberg, L. E., Brookes, D. H., Wilson, L., Chen, J., Liles, K., Chun, M., Li, P., Gohara, D. W., Dolinsky, T., Konecny, R., Koes, D. R., Nielsen, J. E., Head-Gordon, T., Geng, W., Krasny, R., Wei, G. W., Holst, M. J., McCammon, J. A. & Baker, N. A. Improvements to the APBS biomolecular solvation software suite. Protein Sci. 27:112–128 (2018).

39. Jumper, J. et al. Highly accurate protein structure prediction with AlphaFold. Nature 596:583–589 (2021).

40. Watanabe, S., Amagai, Y., Sannino, S., Tempio, T., Anelli, T., Harayama, M., Masui, S., Sorrentino, I., Yamada, M., Sitia, R. & Inaba, K. Zinc regulates ERp44-dependent protein quality control in the early secretory pathway. Nat. Commun. 10:603 (2019).

41. Säbel, E. C., Neureuther, J. M. & Siemann, S. A spectrophotometric method for the determination of zinc, copper, and cobalt ions in metalloproteins using Zincon. Anal. Biochem. 397:218–26 (2010).

42. Parsons, D. S., Hogstrand, C. & Maret, W. The C-terminal cytosolic domain of the human zinc transporter ZnT8 and its diabetes risk variant. FEBS J. 285:1237–1250 (2018).

